# Dynamics-aware geometric learning predicts disease-associated molecular perturbations

**DOI:** 10.64898/2026.08.17.745323

**Authors:** Yuhan Ning, Mingyue Cai, Dongfang Luo, Yuan Li, Gennady Verkhivker, Guang Hu, Zhongjie Liang

## Abstract

Missense mutations and post-translational modifications (PTMs) are major molecular perturbations that reshape protein function but are traditionally studied independently. Current computational approaches largely rely on sequence conservation or static structural features, limiting our understanding of how perturbations alter intrinsic protein dynamics. We present DynGeo-Pheno, a unified geometric deep learning framework that integrates protein language model representations with anisotropic network model–derived dynamics to jointly capture evolutionary, structural, and biophysical information. DynGeo-Pheno predicts disease-associated phosphosites and pathogenic missense mutations with high accuracy on independent test datasets. Ablation analyses indicate that protein dynamics provide complementary information beyond sequence evolution and structural topology for pathogenicity prediction. Beyond predictive performance, DynGeo-Pheno reveals that disease-associated perturbations preferentially localize to functional structural regions, including ligand-binding pockets and PPI interfaces. Mechanistically, phosphosites and missense mutations appear to exhibit distinct yet convergent dynamics signatures. Phosphosites preferentially occur in flexible regulatory regions, whereas pathogenic mutations are enriched in ordered structural elements. Despite these differences, both perturbation types display enhanced long-range coupling, increased perturbation responsiveness, and elevated mechanical stability, indicating that pathogenic residues preferentially occupy mechanically constrained and allosteric regulatory sites. This study provides a compelling evidence that the intrinsic protein dynamics is an important complementary determinant of pathogenicity and establishes a unified framework for interpretable AI predictions and mechanistic understanding of how genetic and regulatory perturbations may shape protein function.

**Significance:** Predicting how genetic mutations and post-translational modifications alter protein function remains a central challenge in human disease research. Most computational approaches focus on sequence or static structural features, while intrinsic protein dynamics remain underutilized. Here, we present DynGeo-Pheno, a unified geometric deep-learning framework that integrates evolutionary information, structural topology, and protein dynamics to model disease-associated phosphosites and pathogenic missense mutations. Although sequence representations capture substantial predictive information, explicit dynamics provides complementary biophysical insights, revealing that pathogenic perturbations preferentially occupy mechanically influential regions characterized by strong dynamics coupling and constrained collective motions. This framework provides a unified strategy for linking local molecular perturbations to the broader structural and dynamical organization of proteins.

## Introduction

Missense mutations and post-translational modifications (PTMs) represent two fundamental layers of molecular variation that shape protein function(1). Missense mutations arise from nucleotide substitutions that alter amino acid sequences, whereas PTMs, such as phosphorylation, are reversible covalent modifications that regulate proteins after translation(2). Both act as perturbations to protein systems, influencing structural stability, biochemical activity, and interaction networks, and are widely implicated in cancer, neurodegeneration, and other human diseases(3). Large-scale proteogenomic studies have further revealed that disease-associated mutations are frequently enriched near PTM sites, highlighting a tight functional coupling between genetic variation and post-translational regulation(4, 5). Accurate characterization of their functional impact is therefore essential for understanding disease mechanisms and developing targeted therapies(6). However, while experimental assays remain the gold standard, their limited scalability has driven the rapid development of computational approaches(7).

For missense mutations, machine learning (ML) and deep learning (DL) methods have achieved substantial progress(8–20), giving rise to a range of high-performing predictors based on sequence evolution (e.g., SIFT(9), PolyPhen-2(10), EVMutation(11)), structural features (e.g., Missense3D(**Error! Reference source not found.**), LYRUS(13)), and integrated functional annotations (e.g., WS-SNPs&GO(14), MutPred2(15)). More recently, embeddings from protein language models (PLMs), such as ESM-2(16), have become foundational inputs, significantly improving predictive performance. For example, ESM-1b enables large-scale evaluation of hundreds of millions of variants using sequence-derived representations(19), while AlphaMissense integrates coevolutionary signals with AlphaFold-based structural embeddings to achieve state-of-the-art pathogenicity prediction(20).

In parallel, extensive efforts have been devoted to phosphosite prediction, spanning general and kinase-specific models, including DeepPhos(21), PhosIDN(22), MusiteDeep(23), GPS 6.0(24), and DeepMVP(25). Advances in representation learning, such as PTM-Mamba(1), further enhance prediction by integrating PTM-aware tokens with PLM embeddings. Structure-based approaches are also evolving with geometric DL frameworks, such as MIND-S(26) and MMFuncPhos(27), incorporating graph neural networks (GNNs) to model local structural environments in PTM and functional phosphosite prediction. Despite these advances, computational models for functional characterization of phosphosites remain largely limited. The representative models combined multi-source features indicative of structural, regulatory, or evolutionary relevance, to make prioritization of functional phosphosites(28–30). Our previously developed DL model, FuncPhos SEQ(31), evaluates the functional potential of phosphosites by integrating sequence motifs and protein–protein interaction information.

A fundamental limitation of current methods is their reliance on sequence evolution and static structural features(32), which inadequately capture the intrinsic conformational dynamics central to protein function(33–35). Both mutations and phosphorylation events can be conceptualized as perturbations to dynamics systems(36). Pathogenic mutations are enriched in regions critical for collective motions, where they modulate intramolecular communication(34, 35). Similarly, phosphorylation primarily reshapes conformational fluctuations, often exerting distal effects without inducing large structural rearrangements(36, 37). These observations underscore the need to incorporate biophysical dynamics into predictive models. Indeed, integrating physics-based representations with ML has proven highly promising(38). For instance, the Rhapsody framework leverages dynamics descriptors derived from elastic network models (ENMs) to improve the prediction of pathogenic mutations(39). Similarly, our previous work, FuncPhos-STR(40), demonstrated that incorporating ENM-derived structural dynamics also enhances the functional prediction of phosphosites.

Despite this progress, current approaches remain largely correlational and descriptive, centering on sequence conservation, static spatial proximity, and limited dynamics features. Such strategies inadequately capture the intrinsic conformational dynamics that underlie protein function, thereby limiting mechanistic insight into how missense mutations and phosphorylation events perturb dynamics-driven regulation. Moreover, existing models typically treat missense mutations and phosphorylation as independent problems, lacking a unified architecture that captures their shared biophysical basis through the integration of systematic dynamics embeddings with geometric GNNs. Meanwhile, resources such as ActiveDriverDB(41), PRISMOID(42), and Missense3D-PTMdb(43) have shown that pathogenic missense variants are frequently enriched near PTM sites in both sequence and 3D space. DL models, including MIND-S(26) and DeepMVP(25), further enable accurate prediction of PTM sites and provide interpretable modules for assessing mutation effects, thereby facilitating the investigation of mutation–PTM relationships. Nevertheless, these approaches still rely predominantly on descriptive analyses based on sequence or structural proximity and fail to resolve the functional coupling between mutation and PTM-mediated conformational dynamics.

Here we present DynGeo-Pheno, a unified and interpretable framework integrating protein dynamics and sequence representations into geometric GNNs. It extracts dynamics embeddings using ANMs(32, 44) and fuses them with ESM-2 embeddings within a multi-scale GCN–GAT architecture, enabling the capture of both protein semantic and structural dynamics information. Furthermore, the framework offers multilevel interpretability, linking predictions to biological function and biophysical mechanisms. Disease-associated phosphosites and mutations are not only enriched in functional regions such as ligand-binding pockets and PPI interfaces(45), but also exhibit distinct dynamics coupling patterns characterized by dynamics cross correlation, perturbation response scanning and mechanical stiffness. By bridging biophysical embeddings with DL, DynGeo-Pheno establishes a dynamics-aware model for the mechanistic interpretation of molecular perturbations, supporting more interpretable predictions and rational therapeutic targeting.

## Results

### Phosphosite/Missense mutation Datasets and Features

We first constructed two curated benchmark datasets spanning phosphorylation and missense mutation phenotypes. The phosphorylation dataset, compiled from PTMD 2.0, PSP, PTMint, and iPTMnet databases and overlapped with PTMAtlas dataset(25, 46–49), contains 3,059 disease-associated phosphosites together with 70,852 neutral sites, mapped to 6,005 human proteins (Table S1). The missense mutation dataset, compiled from ClinVar and overlapped with Rhapsody-2 dataset(50, 51), comprises 89,471 variants across 10,980 proteins, including 30,724 pathogenic and 58,747 neutral missense mutations (Table S2). At the protein level, 3,894 proteins are common to both datasets, with 393 residues showing direct overlap between phosphosites and mutation positions (Fig. 1A). Moreover, neighborhood-level analysis revealed substantially broader structural overlap between mutation-centered and phosphorylation-centered residue neighborhoods than direct site overlap alone, further supporting shared local structural contexts across the two perturbation types (Fig. S1). Characterization of disease annotations (Fig. 1B) shows that the phosphorylation cohort spans diverse disease contexts and exhibits strong enrichment for multiple cancer types, indicating the broad clinical relevance of dysregulated phosphorylation events. In contrast, the missense mutation dataset captures a wider spectrum of inherited and complex disorders, reflecting the heterogeneous phenotypic consequences of coding sequence variation.

**Fig 1.**
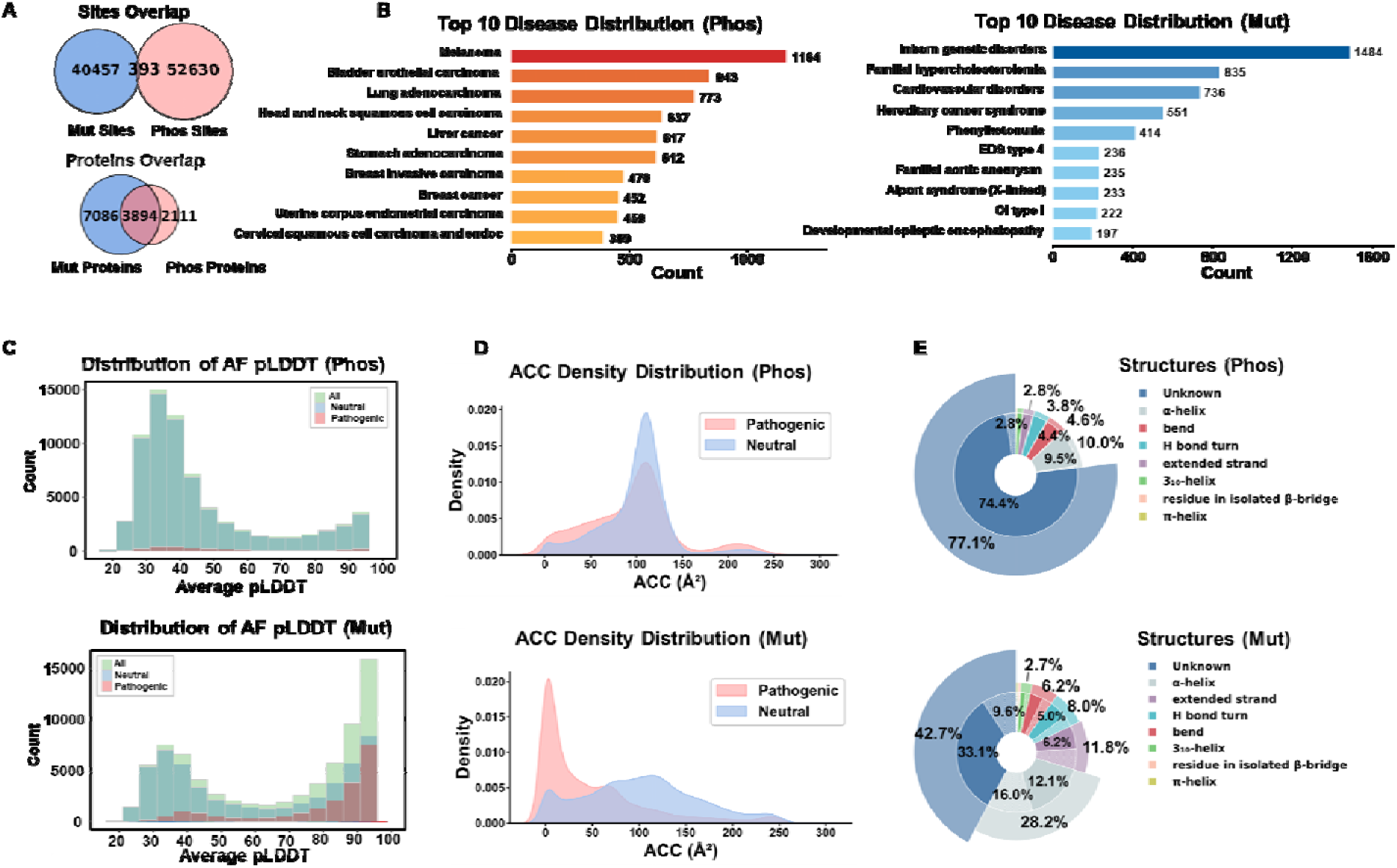
Structural and functional landscape of disease-associated phosphosites and missense mutations. (A) Residue-level and protein-level overlap between phosphosite and mutation datasets. Venn diagrams show shared proteins and direct residue-level overlap between annotated phosphosites and mutation sites. (B) Disease categories associated with curated phosphosites and mutation sites. (C) Distribution of AlphaFold predicted Local Distance Difference Test (pLDDT) for all, neutral, and disease-associated phosphosites and mutation sites. (D) Solvent accessibility (ACC) for phosphosites and mutation sites. (E) Secondary-structure composition of phosphorylation sites and mutation sites.

Mapping all sites onto AlphaFold-predicted structures reveals distinct local structural distributions (Fig. 1C). The pLDDT distributions differ between phosphorylation and mutation sites.

Phosphosites are predominantly located in regions with relatively low confidence (average pLDDT of 30–45), whereas mutation sites exhibit a distinct distribution with an additional peak at high-confidence regions (85–95). Within each dataset, pathogenic and neutral sites show largely overlapping pLDDT distributions. The ACC distributions also differ markedly between phosphorylation and mutation sites (Fig. 1D). Pathogenic and neutral phosphosites show largely overlapping distributions with a common peak around 100 Å². In contrast, pathogenic mutations are strongly enriched at low ACC values, whereas neutral mutations exhibit a broader distribution toward solvent-exposed residues.

Secondary structure analysis (Fig. 1E) shows that phosphosites are predominantly located in residues with unknown secondary structure, accounting for over 70% of all sites, whereas α-helices represent the second most abundant structural class. The structural composition is highly similar between pathogenic and neutral phosphosites. In contrast, mutation sites exhibit a more diverse structural distribution, with a substantially lower proportion of unknown structures and increased enrichment in α-helices and β-strands. Collectively, these observations reveal fundamentally different structural localization patterns between phosphosites and missense mutations, indicating that static structural features capture only part of the underlying functional landscape and underscoring the need to incorporate protein dynamics to better characterize disease-associated perturbations.

### Overview of the DynGeo-Pheno architecture

To jointly model disease-associated phosphosites and pathogenic missense mutations, we developed DynGeo-Pheno, a unified geometric deep learning framework that integrates evolutionary constraints, intrinsic protein dynamics, and three-dimensional structural topology within a common predictive architecture (Fig. 2). Instead of constructing independent models for different classes of molecular perturbations, DynGeo-Pheno adopts a shared geometric learning strategy while preserving task-specific representations of phosphorylation sites and missense mutations.

**Fig. 2.**
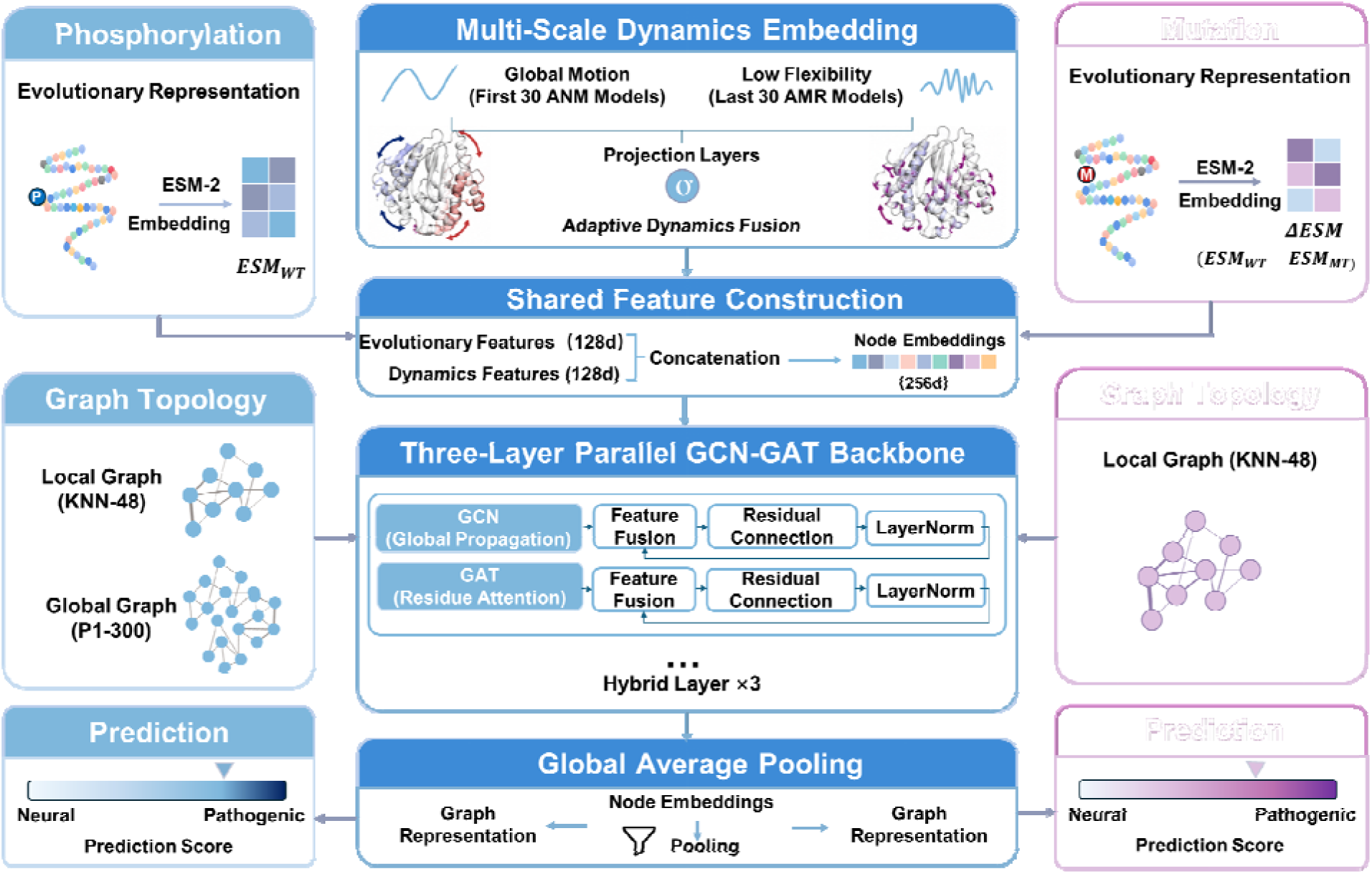
Overview of the DynGeo-Pheno architecture.

The framework is built upon three complementary sources of residue-level information. Evolutionary constraints are captured using ESM-2 protein language model embeddings, where phosphorylation sites are represented by embeddings derived from wild-type sequences, whereas missense mutations are characterized by the difference between mutant and wild-type embeddings (ΔESM) to explicitly encode mutation-induced evolutionary perturbations. To complement sequence-derived information with conformational properties, intrinsic protein dynamics are characterized using the eigenvectors of anisotropic network models (ANMs). Low-frequency collective motions and high-frequency localized fluctuations are jointly integrated through an adaptive gating mechanism, enabling residue representations to capture both global conformational coordination and local mechanical constraints. The resulting dynamics representation is concatenated with evolutionary features to generate residue-level node embeddings. These node embeddings are mapped onto protein structural graphs for subsequent geometric learning. Phosphosite prediction employs both a residue-centered local graph and a protein-level global graph to capture complementary local environments and long-range structural communication, whereas mutation prediction uses a mutation-centered local graph that reflects the localized effects of single amino acid substitutions. All graphs are processed using a shared three-layer parallel GCN–GAT backbone, which combines topology-guided message passing with attention-based neighbor weighting to learn complementary structural representations (Fig. S2).

Finally, graph-level representations are obtained by global average pooling and used for task-specific classification. By integrating evolutionary information, adaptive dynamics representations, and protein structural topology within a unified geometric learning framework, DynGeo-Pheno provides a general computational framework for characterizing residue-level functional vulnerability across multiple classes of disease-associated molecular alterations.

### Benchmarking of DynGeo-Pheno Against State-of-the-Art Datasets

Hyperparameter optimization was performed independently for disease-associated phosphosite prediction and pathogenic missense mutation prediction by systematically evaluating alternative configurations of dynamics representations, evolutionary embedding dimensionality, graph neighborhood size, and structural confidence filtering (Fig. S3, Table S3). Despite the two prediction tasks differing in biological context, both exhibited highly consistent optimization trends. Among the dynamics representations, fusion of the first 30 low-frequency and last 30 high-frequency ANM modes consistently produced the highest AUROC, indicating that global collective motions and local mechanical constraints provide complementary information for residue-level functional prediction. This further supports our central hypothesis that residue functional vulnerability is governed by complementary conformational motions operating across multiple spatial scales. Similarly, reducing ESM embeddings to 128 dimensions achieved the best predictive performance, suggesting that this representation effectively preserves informative evolutionary signals while avoiding redundant features.

Model performance was also sensitive to graph construction parameters. A neighborhood size of K = 48 yielded the highest AUROC for both tasks, supporting a balance between capturing sufficient structural context and minimizing the introduction of irrelevant neighboring residues. In contrast, applying increasingly stringent pLDDT confidence thresholds consistently reduced prediction performance, whereas retaining all residues without confidence filtering achieved the best results. This observation suggests that structurally less confident regions, which frequently correspond to flexible or intrinsically dynamics segments, still contain functionally relevant information for identifying disease-associated phosphosites and pathogenic missense mutations. Based on these analyses, the final DynGeo-Pheno model adopted the fusion of the first 30 low- and last 30 high-frequency ANM modes, 128-dimensional ESM embeddings, K = 48, and no pLDDT filtering for all subsequent experiments.

To evaluate the generalizability of DynGeo-Pheno, we first trained the framework using ten-fold cross-validation on the training and validation datasets and subsequently evaluated the resulting ensemble models on independent test datasets for disease-associated phosphosite and pathogenic missense mutation prediction. DynGeo-Pheno achieved consistently high predictive performance on the independent test datasets, yielding AUROC values of 0.873 for disease-associated phosphosite prediction and 0.937 for pathogenic mutation prediction, together with AUPRC values of 0.772 and 0.896, respectively (Fig. 3A, B; Table S4). The ROC curves obtained from individual cross-validation folds showed minimal variation, indicating that the framework learned stable representations with good generalization ability across independent data partitions. We next compared DynGeo-Pheno with representative state-of-the-art predictors using multiple evaluation metrics, including accuracy, precision, recall, F1 score, AUROC, AUPRC, and MCC. (Fig. 3C). For disease-associated phosphosite prediction, DynGeo-Pheno consistently outperformed FuncPhos-SEQ and FuncPhos-STR across most evaluation metrics. AUROC increased from 0.692 and 0.753 to 0.864, respectively, while MCC largely improved from 0.265 and 0.361 to 0.572. For pathogenic missense mutation prediction, DynGeo-Pheno achieved the highest recall (0.885), while maintaining AUROC (0.914) and AUPRC (0.880) comparable to those of AlphaMissense and consistently higher than those of Rhapsody-2. Given differences in modeling objectives and input representations, we consider AlphaMissense a complementary reference rather than a direct competitor. This comparison not only provides external validation for DynGeo-Pheno but also underscores the unique value of integrating structural topology and intrinsic protein dynamics for pathogenicity prediction. These results demonstrate that integrating evolutionary information and protein dynamics with geometric graph learning achieves performance comparable to large protein language models while using a substantially more compact architecture.

**Fig. 3.**
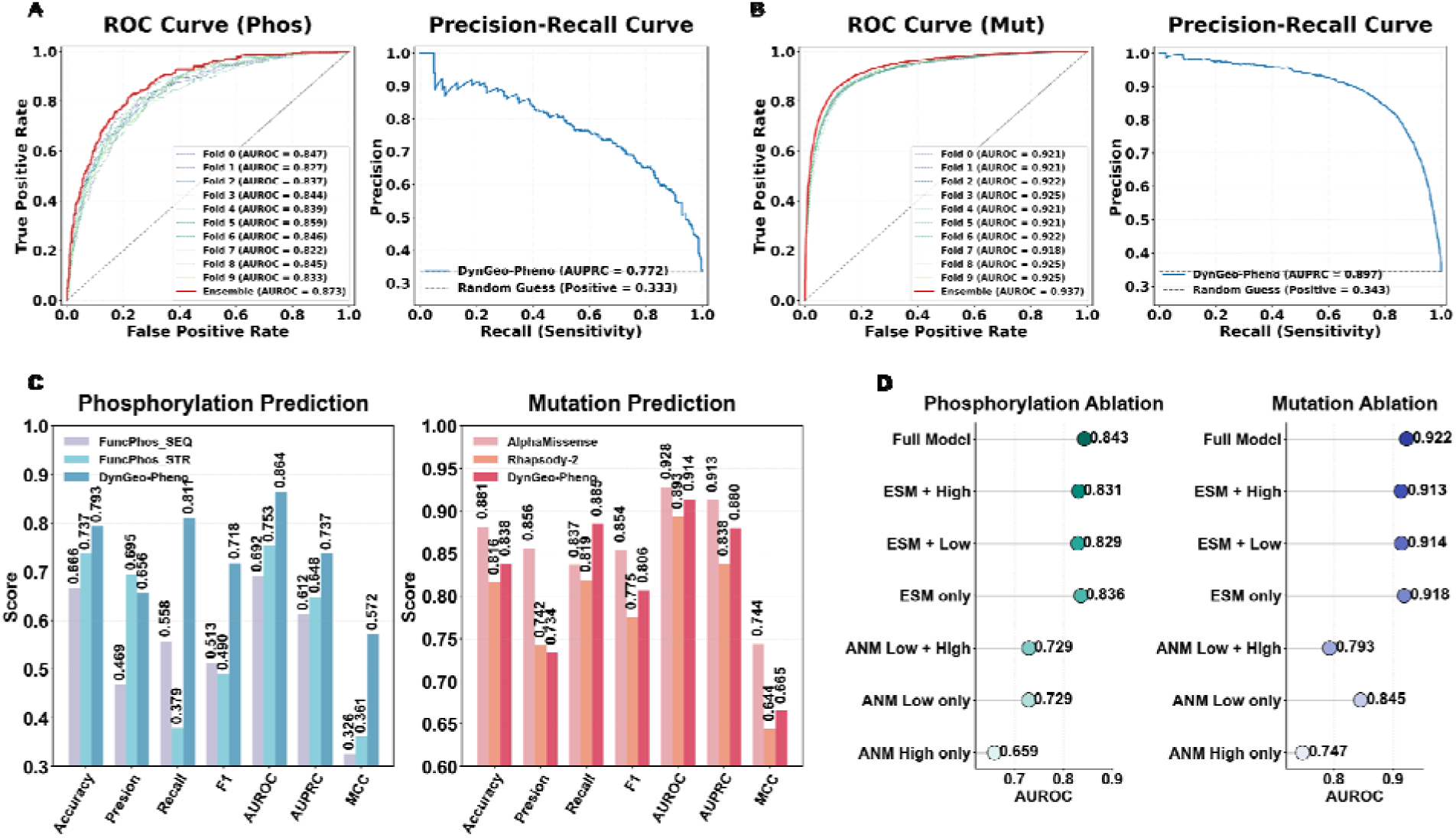
Benchmark evaluation and feature contribution analysis of DynGeo-Pheno. (A) Receiver operating characteristic (ROC) and precision–recall (PR) curves for disease-associated phosphosite prediction. Insets show ROC curves from the ten cross-validation folds used to construct the ensemble model. (B) ROC and PR curves for pathogenic missense mutation prediction, with fold-specific ROC curves shown in the inset. (C) Performance comparison of DynGeo-Pheno with representative state-of-the-art methods for disease-associated phosphosite prediction (left) and pathogenic missense mutation prediction (right). Model performance was evaluated using accuracy, precision, recall, F1 score, AUROC, AUPRC, and Matthews correlation coefficient (MCC). (D) Feature ablation analysis showing the contributions of ESM-derived evolutionary representations and ANM-derived dynamics features. AUROC values are reported for individual feature modalities and their combinations in phosphosite (left) and missense mutation (right) prediction.

### Ablation Study

To evaluate the contribution of each information modality, we performed systematic feature ablation analyses by removing evolutionary or dynamics representations (Fig. 3D and Table S5). Across both prediction tasks, ESM-2 embeddings alone provided strong predictive performance, highlighting the importance of sequence-derived functional constraints. Nevertheless, incorporating ANM-derived dynamics representations further improved the full model, which achieved the highest AUROC for both phosphosite and missense mutation prediction. The Full Model also showed improved AUPRC and recall values (Table S5), indicating enhanced identification of disease-associated residues, an advantage that is particularly relevant to imbalanced datasets and may not be fully captured by accuracy alone. The relative contributions of low- and high-frequency dynamics modes varied between the two prediction tasks. However, their adaptive integration in the Full Model consistently outperformed models using either frequency component alone, suggesting that global collective motions and local mechanical fluctuations encode complementary, task-dependent information. These results support the value of integrating multiple dynamics representations with evolutionary features rather than relying on any single information modality. While sequence-derived representations capture the majority of predictive signal, explicit modeling of protein dynamics provides consistent, statistically significant complementary information and, more importantly, reveals the biophysical mechanisms underlying pathogenicity.

### Biological and dynamics interpretation of DynGeo-Pheno predictions

To verify whether predictions from DynGeo-Pheno capture functionally meaningful patterns rather than purely correlative signals, we systematically evaluated model outputs from both biological and dynamics perspectives (Fig. 4). These complementary analyses were conducted within a unified three levels of evaluation. First, we examined whether disease-associated perturbations exhibit distinct higher prediction scores relative to neutral ones, both inside and outside biological or dynamics “hot” regions. Second, we quantified the preferential association between perturbed residues and feature localization using the odds ratio (OR), which measures the relative likelihood of observing a disease-associated residue inside a given region compared to neutral residue. Third, as the OR does not account for regional size or residue density, we further computed positive and negative density ratios (Positive DR and Negative DR), which respectively quantify the relative density enrichment of disease-associated and neutral residues within each region, normalized against the protein-wide background. Together, these metrics provide complementary assessments of preferential localization, hot region association, and hot region enrichment.

**Fig. 4.**
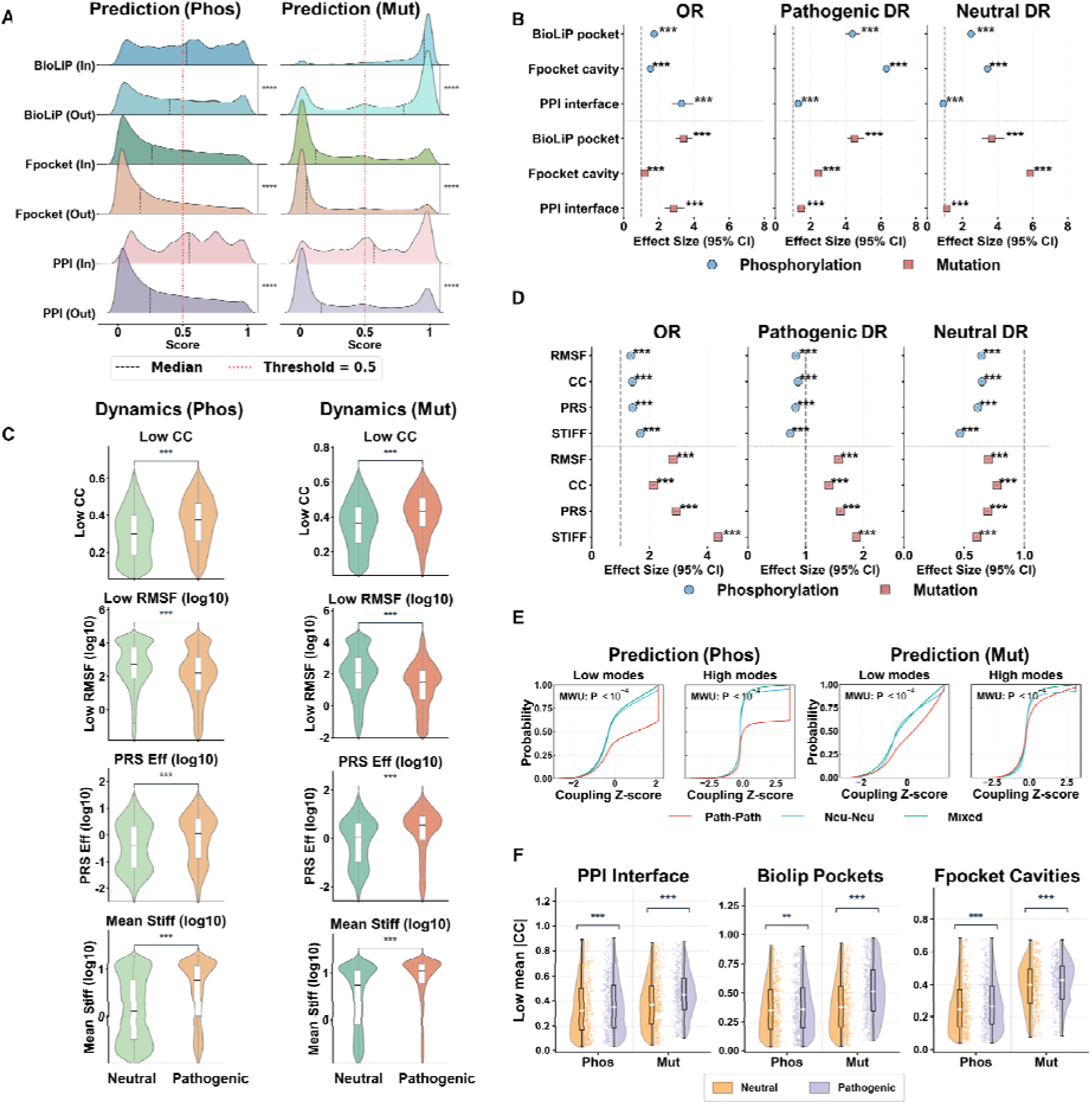
Biological and dynamics interpretation of DynGeo-Pheno predictions. (A) Prediction score distributions for disease-associated and neutral residues located inside and outside annotated functional regions, including BioLiP ligand-binding pockets, Fpocket cavities, and protein–protein interaction (PPI) interfaces, for phosphosites (left) and missense mutations (right). The dashed lines indicate the optimal threshold, defined as the median. (B) Association and enrichment of disease-associated residues in annotated functional regions. Odds ratio (OR), positive density ratio (Positive DR), and negative density ratio (Negative DR) are shown for phosphosites (blue) and missense mutations (red) in BioLiP pockets, Fpocket cavities, and PPI interfaces. Error bars indicate 95% confidence intervals. Statistical significance was assessed using two-sided Fisher’s exact tests with Benjamini–Hochberg correction. (C) Comparison of representative ANM-derived dynamics descriptors between disease-associated and neutral residues, including residue cross-correlation (CC), root mean square fluctuation (RMSF), perturbation response scanning (PRS) effectiveness, and mechanical stiffness. (D) Association and enrichment of disease-associated phosphosites and mutations within dynamics hotspot regions defined by CC, RMSF, PRS effectiveness, and stiffness, quantified by OR, Positive DR, and Negative DR. (E) Distribution of ANM coupling scores for disease-associated and neutral residue pairs based on low- and high-frequency modes for phosphosites (left) and mutations (right). (F) ANM coupling scores between disease-associated residues and annotated functional regions, including PPI interfaces, BioLiP pockets, and Fpocket cavities, after excluding residues directly located within the corresponding regions. Statistical significance for Figure 4(A), 4(C) and 4(E-F) was assessed using Mann-Whitney U test.

Disease-associated residues consistently yielded higher prediction scores than neutral residues within experimentally validated functional regions, including BioLiP pockets (Table S6, S7)(52) and PPI interfaces (Table S8, S9), whereas this separation was markedly weaker within Fpocket predicted cavities (Table S10, S11) (Fig. 4A)(53). Consistent with these observations, OR analysis revealed these disease-associated residues are strongly associated with pocket regions and PPI interfaces. Moreover, enrichment analysis revealed significant overrepresentation of disease-associated residues in BioLiP pockets and PPI interfaces for both prediction tasks, whereas the enrichment within purely geometric cavities was comparatively modest (Fig. 4B). Collectively, these findings indicate that DynGeo-Pheno preferentially identifies biologically functional regions rather than merely structurally accessible regions.

We next investigated whether the model preferentially detects disease-associated residues with characteristic dynamics properties, including (i) RMSF, where minima indicate the sites that potentially act as hinges for supporting protein’s mechanics; (ii) cross-correlation, which indicates the dynamics coupling between residue pairs; (iii) the propensity of residues to act as sensors or effectors of allosteric signals based on perturbation-response scanning (PRS) analysis; (iv) the mechanical stiffness associated with each residue. As shown in Fig. 4C, disease-associated residues consistently exhibited higher cross-correlation, greater PRS effectiveness, increased mechanical stiffness, and lower RMSF than neutral residues (Table S12, S13), with additional ENM-derived descriptors exhibiting consistent patterns (Fig. S4), indicating that these residues preferentially reside in mechanically constrained and communication-active regions. Building on these observations, residues falling within the extreme 20% of each ANM-derived descriptor (lowest RMSF; highest CC, PRS effectiveness, and stiffness) were defined as intrinsic dynamics hotspots. OR analysis revealed these disease-associated residues are strongly associated with the dynamics hot regions. Enrichment analyses demonstrated significant overrepresentation of disease-associated residues across all four hotspot categories, accompanied by elevated positive DR, and reduced negative DR values (Fig. 4D). These results suggest that disease-associated perturbations preferentially occur at structurally constrained and dynamically influential positions, rather than being randomly distributed throughout protein structures.

Finally, we examined whether the functional influence of disease-associated residues extends beyond their local structural contexts. Pairwise dynamics cross correlation analyses revealed significantly stronger dynamics coupling between disease-associated residue pairs than between neutral or mixed residue pairs, indicating that disease-associated sites preferentially participate in coordinated communication networks (Fig. 4E). Importantly, even after excluding residues directly located within BioLiP pockets, Fpocket cavities, and PPI interfaces, disease-associated residues remained significantly more strongly coupled to functional regions than neutral residues (Fig. 4F). Similar coupling patterns were observed when the coupling was evaluated using high-frequency ANM modes (Fig. S5). These observations suggest that disease-associated perturbations propagate through intrinsic communication pathways rather than being restricted to experimentally annotated functional sites, providing a mechanistic basis for the functional impact of distal residues.

### Case study

GO and KEGG analyses of proteins harboring structurally enriched disease-associated residues revealed distinct functional signatures (Fig. S6). Proteins with pocket-localized phosphosites were enriched in kinase signaling, small GTPase regulation, and cytoskeletal organization, reflecting the regulatory role of phosphorylation in dynamics signaling networks. Conversely, pocket-localized pathogenic mutations preferentially occurred in proteins involved in ATP-dependent catalysis, conformational regulation, and metabolic pathways, consistent with disruption of constrained functional cores. At PPI interfaces, phosphosite-enriched proteins were associated with signaling and immune regulation, whereas mutation-enriched proteins were linked to receptor interactions, apoptosis, and oncogenic processes. These results demonstrate that DynGeo-Pheno identifies structurally and functionally distinct disease-associated modules associated with different molecular perturbation mechanisms.

To further evaluate the ability of DynGeo-Pheno to identify pathogenic perturbations, we performed in-depth analyses of two representative systems, BRAF, and MTHFR. BRAF is a central kinase in the MAPK/ERK signaling pathway that regulates cell proliferation and survival(54). Mapping predicted perturbation sites onto the BRAF structure revealed distinct spatial distributions of pathogenic and neutral residues (Fig. 5A). Among the identified sites, 12 positive and 8 negative mutations, as well as 6 positive and 8 negative phosphosites, were characterized. Positive mutations are preferentially localized to the structurally constrained core, whereas positive phosphosites were enriched in regulatory regions. Notably, many positive sites overlapped with ligand-binding pocket(55) and PPI interface(56). In contrast, negative sites were sparsely scattered within flexible loop regions. Consistent with their predicted pathogenicity, positive sites—including mutations G464E/R/V, G469A/E/R/V, and D594N/V, as well as phosphorylation events T599-p and S614-p—have been functionally characterized in previous studies(57–59). whereas no functional evidence has been reported for the negative sites. We next examined MTHFR, a key metabolic enzyme involved in folate metabolism, DNA synthesis and epigenetic regulation(60). Similarly, predicted positive sites (5 mutations and 1 phosphosite) were preferentially localized within ligand-binding pockets(61), whereas negative sites showed no apparent structural enrichment (Fig. 5B). Several positive mutations, including R157Q, A175T, A195V, and P572L, have been experimentally linked to MTHFR dysfunction (62, 63). Additional analyses of TAOK1 and KCNH1 further supported this observation, as positive sites in both systems showed significant overlap with Fpocket-predicted pockets (Fig. S7A and B).

**Fig. 5.**
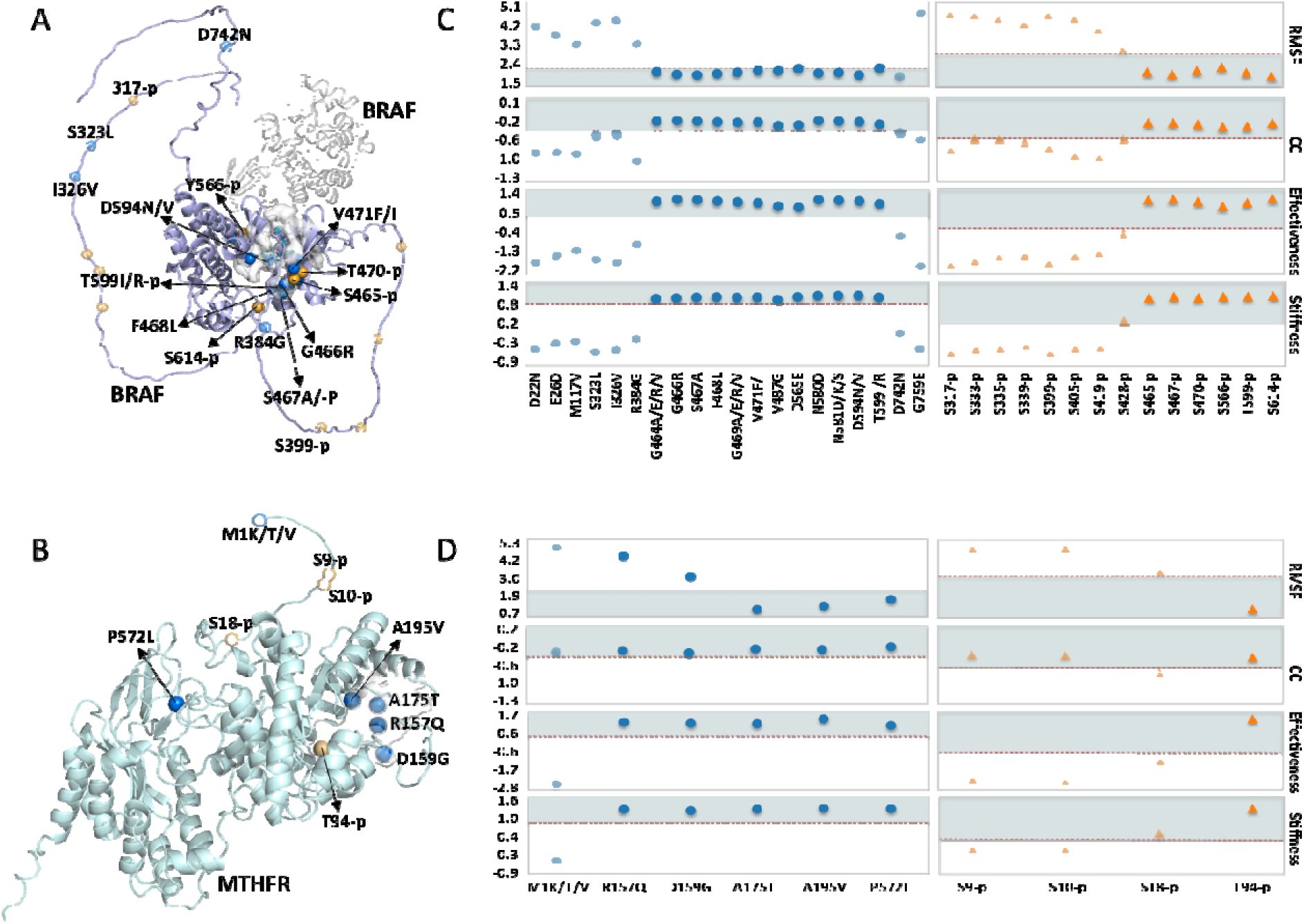
Representative case studies. (A, B) Structural representations of BRAF (A) and MTHFR (B). BRAF is colored purple (with certain long chains omitted for clarity), accompanied by its interacting partner in gray (PDB ID:1UWH). MTHFR is colored cyan. Ligand-binding pockets (PDB IDs: 8DGT and 4FC0 for BRAF; 6FCX for MTHFR) are depicted as white surfaces. Mutations and phosphosites are represented by blue and orange spheres, respectively. Positive sites are displayed at full opacity. Negative sites are rendered with 50% transparency. (C, D) Scatter plots illustrating the four evaluation metrics for the BRAF (C) and MTHFR (D) systems. The dashed red lines indicate the optimal thresholds, defined as the medians. Shaded areas represent the qualified regions for positive sites. Mutations and phosphosites are designated by blue circles and orange triangles, respectively. Opacity standards remain consistent with the structural representations.

Across these representative systems, disease-associated perturbations consistently exceeded the predefined thresholds for all four evaluation metrics, whereas predicted neutral sites largely remained outside the qualified regions (Fig. 5 C and D, Fig. S7C and D), demonstrating clear discrimination between pathogenic and neutral perturbations. Despite substantial differences in sequence and structural organization, these proteins exhibited conserved spatial patterns for both mutations and phosphorylation events, suggesting that DynGeo-Pheno captures shared structural and dynamics determinants underlying diverse molecular perturbations. Together, these findings demonstrate that integrating intrinsic protein dynamics with geometric structural representations enables accurate identification and mechanistic interpretation of functionally important perturbation sites.

## Discussion and Conclusion

### Dynamics as complementary information in pathogenicity prediction

A central premise underlying most existing pathogenicity predictors is that sequence conservation and static structural context provide sufficient information to infer the functional consequences of genetic variants(8, 16). However, proteins function as dynamics systems in which biological activity emerges from coordinated conformational fluctuations and long-range allosteric communication rather than static three-dimensional conformations alone(64, 65). Consequently, functional perturbations caused by disease-associated phosphorylation events or missense mutations are often manifested through alterations of intrinsic protein dynamics, a dimension that remains underrepresented in current predictive frameworks(36).

Our results show that protein language model embeddings coupled with geometric deep learning account for the majority of predictive performance, whereas dynamics embeddings derived from both low-frequency collective motions and high-frequency local vibrational modes provide consistent, albeit modest, improvements in prediction accuracy (0.7% AUROC for phosphosites and 0.4% for missense mutations). The relatively small gain is consistent with the hierarchical relationship in which sequence determines structure and structure constrains intrinsic dynamics, implying that evolutionary representations already encode substantial structural and dynamics information(66–68). This hierarchy likely underlies the strong performance of sequence-based predictors such as AlphaMissense(20), while the success of Rhapsody-2 further demonstrates the predictive value of explicitly incorporating protein dynamics(50). The consistent improvement achieved by explicitly modeling dynamics indicates that protein motions contain complementary information beyond evolutionary representations(34). Whereas sequence-derived embeddings primarily capture evolutionary constraints(20), dynamics directly characterizes residue flexibility, mechanical coupling, and collective motions that determine how local perturbations propagate through protein structures. These observations suggest that protein dynamics may provide an additional mechanistic layer linking molecular variation to phenotypic outcomes and offers a biophysical rationale for integrating dynamics with sequence-derived representations in geometric deep learning framework.

### Dynamics reveals distinct biophysical mechanisms underlying disease-associated phosphosites and pathogenic mutations

Beyond improving prediction accuracy, a principal contribution of this study is the identification of distinct structural and dynamics principles underlying disease-associated phosphosites and pathogenic missense mutations. Although these two classes of pathogenic residues occupy different structural environments, they exhibit convergent dynamics properties that are not apparent from sequence or static structures alone. Consistent with previous studies, disease-associated phosphosites are preferentially enriched within intrinsically disordered regions and flexible loops characterized by relatively low AlphaFold confidence (pLDDT) and poorly defined secondary structures(29–32). Such environments facilitate kinase recognition, reversible modification, and context-dependent regulation. In contrast, pathogenic missense mutations occur predominantly within ordered α-helices and β-strands with high structural confidence, where amino acid substitutions are more likely to disrupt residue packing, hydrogen-bonding networks, and structural stability(12, 15).

Despite these contrasting structural preferences, both classes of pathogenic residues display remarkably similar dynamics signatures relative to neutral sites, pathogenic phosphosites and missense mutations exhibit stronger low-frequency residue cross-correlations, reduced fluctuations in global collective modes, increased fluctuations in high-frequency local modes, higher perturbation effectiveness, and greater mechanical stiffness. These features indicate that pathogenic residues preferentially occupy mechanically influential positions capable of efficiently transmitting conformational perturbations through residue interaction networks. Interestingly, phosphosites located within structurally flexible regions are dynamically constrained in global motions while remaining highly active in local motions. Their combination of enhanced collective communication, reduced global flexibility, elevated local fluctuations, and increased stiffness suggests that they function as regulatory hotspots that coordinate long-range allosteric signaling while preserving local adaptability required for reversible phosphorylation(64, 69). By contrast, pathogenic missense mutations are concentrated within mechanically rigid structural cores, where local perturbations are more likely to destabilize the mechanical framework and propagate structural defects throughout the protein(70). Together, these observations suggest that phosphorylation may more frequently rewire regulatory communication, whereas pathogenic mutations may more commonly compromise mechanical stability, yet both appear to ultimately perturb the dynamics networks governing protein function.

These mechanistic insights also explain why dynamics improves pathogenicity prediction. Low-frequency modes describe cooperative conformational transitions that mediate long-range allosteric communication, whereas high-frequency modes capture local rigidity and mechanical constraints that stabilize protein architecture(71, 72). Their adaptive integration therefore provides complementary multiscale information that cannot be fully recovered from sequence conservation or static structures alone, supporting the view that disease susceptibility is fundamentally a multiscale dynamics property(73).

### Limitations and Future directions

Several limitations should be acknowledged. First, intrinsic dynamics were estimated from a single reference structure using elastic network models and therefore do not capture large conformational transitions or the full ensemble of accessible protein states(73, 74). Second, the current graph representation focuses primarily on monomeric structures and does not explicitly model inter-subunit communication or cooperative dynamics within multimeric protein complexes(67, 68). In addition, disease-associated phosphosites and pathogenic missense mutations were modeled independently, although these perturbations frequently coexist and interact in vivo(4, 5). Future models should explicitly consider their combined effects to better characterize disease mechanisms. Likewise, our framework currently integrates sequence, structure, and intrinsic dynamics but does not incorporate context-specific molecular information such as quantitative phosphoproteomics, mutation burden, or cell-type-specific gene expression, all of which provide physiological context beyond molecular structure(6, 7).

Looking forward, integrating evolutionary, structural, dynamics, and multi-omics information within unified multimodal learning frameworks may provide a more complete understanding of how molecular perturbations collectively reshape protein function and cellular phenotypes(50). Such approaches could improve the identification of disease-driving events and establish a general framework linking molecular biophysics with systems-level biology, thereby advancing mechanism-informed precision medicine(35).

## Material and Methods

### Framework Overview

To evaluate the hypothesis that disease-associated molecular events preferentially localize to dynamically constrained yet functionally communicative positions within protein structures, we developed DynGeo-Pheno, a multimodal framework that integrates intrinsic protein dynamics, evolutionary sequence information, and geometric structural features for residue-level phenotype prediction (38).

### Dataset Curation and Structural Mapping

We constructed two benchmark datasets for disease-associated phosphorylation site prediction and pathogenic missense mutation prediction. The phosphorylation dataset, compiled from the PTMD 2.0, PSP, and iPTMnet databases and intersected with high-confidence mass spectrometry data from PTMAtlas(28, 29), comprises 73,911 phosphosites across 6,005 human proteins, including 3,059 disease-associated phosphosites and 70,852 neutral phosphosites without functional annotation. The missense mutation dataset, compiled from ClinVar and intersected with the Rhapsody-2 dataset(50), comprises 89,471 curated and standardized variants across 10,980 human proteins, including 30,724 pathogenic and 58,747 neutral variants. Three-dimensional protein structures were retrieved from the AlphaFold Protein Structure Database(75), and all residues were mapped onto their corresponding predicted structures. The distributions of pLDDT confidence scores were analyzed to characterize the structural confidence of phosphorylation and mutation sites and to evaluate the contribution of structural confidence to model performance.

Each dataset was randomly divided into a training set (90%) and an independent test set (10%). To control class imbalance and maintain a consistent class ratio across evaluation stages, stratified random undersampling was subsequently performed independently within the training and independent test sets for phosphosite data. The resulting datasets contained disease-associated and neutral phosphosites at a positive-to-negative ratio of 1:2. In contrast, the missense mutation dataset exhibited an approximately balanced class distribution (∼1:1.9) and therefore no additional resampling was performed. Model development, hyperparameter optimization, and performance estimation were performed exclusively on the training set using 10-fold stratified cross-validation, whereas the held-out test set was used only for final model evaluation. Detailed procedures for data preprocessing, sequence redundancy reduction using CD-HIT (30% sequence identity threshold), and benchmark dataset construction are provided in the SI Appendix(76).

### Multi-modal Representation Learning and Adaptive Feature Fusion

Evolutionary sequence representations were generated using the pre-trained ESM-2 protein language model(16). For disease-associated phosphosite prediction, residue embeddings were extracted directly from the wild-type protein sequence. For missense mutation prediction, wild-type and mutant sequences were encoded independently, and the difference between their residue embeddings (WT − Mut) was used to represent mutation-induced evolutionary perturbations. The original 1,280-dimensional embeddings were projected to 128 dimensions by principal component analysis (PCA) before graph construction. Residue-level dynamic features were computed from anisotropic network models (ANMs) constructed from AlphaFold-predicted structures with a 15.0 Å interaction cutoff, using ProDy(77, 78). Eigenvectors obtained from spectral decomposition of the ANM Hessian matrix were used to characterize intrinsic protein motions. For each selected normal mode, the three Cartesian components corresponding to each residue were extracted and concatenated to generate residue-level dynamics embeddings. After excluding the six trivial rigid-body modes, the first 30 nontrivial low-frequency modes and the last 30 high-frequency modes were encoded independently, yielding two complementary 90-dimensional feature vectors that describe collective and localized motions, respectively.

Low-frequency collective motions and high-frequency localized fluctuations were integrated through a learnable gating module. The gating weights were optimized jointly with the entire network during end-to-end training, allowing the model to adaptively balance the contributions of the two dynamical components for each residue. This dynamics embedding was subsequently combined with the task-specific evolutionary sequence representation to define the node attributes of the residue graph for downstream geometric graph learning(16, 38).

### Geometric Deep Learning Architecture

Proteins were represented as undirected residue graphs, in which edges were defined between residue pairs with Cα–Cα distances of less than 10 Å. The prediction network consisted of a three-layer hybrid architecture combining graph convolutional network (GCN)(79) and multi-head graph attention network (GAT) layers(80). GCN layers aggregated information from neighboring residues to encode local structural topology, whereas GAT layers assigned attention weights to neighboring nodes to model heterogeneous residue interactions. Multi-modal residue features were propagated through parallel GCN and GAT branches with residual connections, followed by graph-level pooling and a multilayer perceptron for residue-level classification (Fig. S2).

### Model Training and Unified Evaluation

To ensure robust generalization performance, we implemented a 10-fold stratified cross-validation strategy with RMSprop optimization. For test set evaluation, final binary classification outcomes for each sample were determined by majority voting across the ten independently trained models. An identical training and evaluation pipeline was applied to both prediction tasks to enable consistent assessment of generalizability. Performance was rigorously assessed using threshold-independent metrics, including AUROC and the area under the precision--recall curve (AUPRC). The latter metric is particularly critical given the pronounced class imbalance of the phosphorylation dataset. Additional classification metrics included the Matthews correlation coefficient (MCC) and F1 score.

### Biological and Dynamics Interpretation Analyses

To evaluate the functional relevance of DynGeo-Pheno predictions, experimentally annotated ligand-binding sites, predicted protein cavities, and PPI interfaces were mapped onto protein structures for post hoc interpretation. In parallel, ANM-derived descriptors, including root mean square fluctuation (RMSF), cross-correlation (CC), perturbation response scanning (PRS) effectiveness(81), and mechanical stiffness(32, 67), were computed to characterize intrinsic protein dynamics. For each descriptor, residues ranked within the top 20% of the protein-specific distribution were defined as dynamics “hotspots” regions and used for enrichment analyses based on odds ratios (ORs) and density ratios (DRs). Gene Ontology (GO) and Kyoto Encyclopedia of Genes and Genomes (KEGG) enrichment analyses were subsequently performed using proteins containing disease-associated residues that showed significant structural enrichment within ligand-binding pockets or PPI interfaces (Positive DR > 1 and adjusted *P* < 0.05), thereby providing a functional interpretation of structurally enriched disease-associated regions(82, 83). Detailed annotation procedures and mathematical definitions are provided in the SI Appendix.

## Statistical Analysis

Differences between continuous variables were evaluated using two-sided Mann–Whitney U tests, while Fisher’s exact tests were employed for categorical enrichment analyses. Multiple hypothesis testing was controlled using the Benjamini–Hochberg false discovery rate (FDR) correction(84). Confidence intervals for enrichment statistics were estimated using 1,000 bootstrap iterations where appropriate. Statistical significance was defined as adjusted P < 0.05. Comprehensive mathematical formulations of the ANM descriptors, GNN layers, and gating operations are provided in the SI Appendix. Detailed model hyperparameter specifications are also provided therein.

## Code availability

The source code used in this study is publicly available at https://github.com/ComputeSuda/DynGeo-Pheno.

## Acknowledgements

This work was also supported by the National Natural Science Foundation of China (NSFC) (22377089 to Z.L., and 32271292 to G.H.) and Project of Biomedical Basic Research Center (BBRC) of Jiangsu, Soochow University. This work was supported by the Priority Academic Program Development of Jiangsu Higher Education Institutions (PAPD). This work was also supported by the National Institutes of Health under Award 1R01AI181600-01 and Subaward 6069-SC24-11 to G.V.

