## Supporting Materials and Methods, Supporting Figures S1-S7, Supporting Tables S1-S13 for "Dynamics-aware geometric learning predicts disease-associated molecular perturbations": SUPPORTING_INFORMATION.docx

Yuhan Ning^1#^, Mingyue Cai^1#^, Dongfang Luo^1^, Yuan Li^1^, [Gennady Verkhivker](https://pubmed.ncbi.nlm.nih.gov/?term=Verkhivker+G&cauthor_id=40721360)^2,3,4*^, Guang Hu^1^, and Zhongjie Liang^1*^

^1^Biomedical Basic Research Center (BBRC) of Jiangsu, School of Life Sciences, Suzhou Medical College of Soochow University, Suzhou 215123, Jiangsu, China

^2^Keck Center for Science and Engineering, Schmid College of Science and Technology, Chapman

University One University Drive, Orange, CA 92866, United States of America

^3^Department of Biomedical and Pharmaceutical Sciences, Chapman University School of

Pharmacy, Irvine, CA 92618, United States of America

^4^Department of Pharmacology, Skaggs School of Pharmacy and Pharmaceutical Sciences,

University of California San Diego, 9500 Gilman Drive, La Jolla, CA 92093, United States of America

\# These authors contributed equally

### Supplementary Materials and Methods

#### 1. Data Collection and Quality Control Protocols

##### 1.1 Construction of Phosphorylation Datasets

Phosphorylation data was collected from four curated resources, including PTMD 2.0, PhosphoSitePlus (PSP), PTMint, and iPTMnet(1-4). A total of 128,660 experimentally supported serine, threonine, and tyrosine phosphorylation sites were obtained after removing duplicate records. Phosphosites explicitly annotated as disease-associated in the source databases were assigned as positive samples (label = 1). Sites without any functional annotations were initially considered candidate negative samples. To reduce the inclusion of uncharacterized phosphosites in the negative set, candidate negatives were intersected with 194,903 high-confidence phosphosites from PTMAtlas(5), which are supported by stringent mass spectrometry quality-control criteria. Only phosphosites satisfying the PTMAtlas confidence criteria were retained as negative samples. To reduce sequence redundancy, protein sequences were clustered using PSI-CD-HIT with a sequence identity threshold of 30%, and a single representative sequence was retained from each cluster(6). The final benchmark dataset (Table S1) contained 73,911 phosphorylation sites, including 3,059 disease-associated sites and 70,852 neutral sites, mapped to 6,005 human proteins.

##### 1.2 Construction of Missense Mutation Datasets

Missense mutations were curated from the ClinVar database (GRCh38 assembly)(7) to construct a benchmark dataset for pathogenicity prediction. Mutations annotated as "Pathogenic" or "Likely pathogenic" were assigned as positive samples (label = 1), whereas mutations annotated as "Benign" or "Likely benign" were assigned as negative samples (label = 0). Mutations of uncertain significance (VUS) were excluded from all analyses. To assess the consistency of clinical annotations, the curated dataset was compared with the Rhapsody-2 benchmark dataset. Among 100,109 overlapping mutations, 99.98% showed identical clinical classifications, indicating high concordance between the two datasets. Protein sequences were clustered using CD-HIT with a 30% sequence identity threshold to reduce sequence redundancy(6), and one representative sequence was retained from each cluster. The final benchmark dataset (Table S2) comprised 89,471 nonredundant missense mutations from 10,980 human proteins, including 30,724 pathogenic and 58,747 neutral mutations.

##### 1.3 AlphaFold Structural Curation

Three-dimensional protein structures were retrieved from the AlphaFold Protein Structure Database and used for graph construction and protein dynamics analysis(8, 9). Proteins longer than 2,700 amino acid residues were excluded to ensure computational tractability. The distributions of predicted Local Distance Difference Test (pLDDT) scores were subsequently compared between positive and negative samples for both prediction tasks to assess the potential contribution of structural confidence to model performance (Fig. 1C).

#### 2. Multi-modal Feature Encoding

##### 2.1 Sequence Semantic Embedding

Protein sequences were encoded using the pre-trained ESM-2 language model (esm2_t33_650M_UR50D)(10), which generates contextualized residue representations capable of capturing evolutionary and biochemical constraints. For a protein sequence comprising $N$ residues, ESM-2 generated a residue-level embedding matrix $E\in\mathbb{R}^{N\times1280}$, where each residue was represented by a 1,280-dimensional feature vector. To reduce feature redundancy and computational complexity, principal component analysis (PCA) was applied to project the original high-dimensional embeddings into a compact 128-dimensional latent space, yielding $E_{\text{PCA}}\in\mathbb{R}^{N\times128}$. To strictly prevent data leakage, the PCA projection matrix was fitted exclusively on the training dataset and subsequently applied to the validation and test sets.

For missense mutation prediction, mutation-induced sequence perturbations were represented by the difference between the PCA-reduced embeddings of the wild-type and mutant residues:

$$\begin{aligned} \Delta E=E_{\text{WT}}-E_{\text{Mut}}\#\left（ 1 \right） \end{aligned}$$

where $E_{\text{WT}}$ and $E_{\text{Mut}}\in\mathbb{R}^{128}$ represent the PCA-reduced embeddings of the wild-type and mutant residues, respectively. For phosphorylation prediction, only wild-type embeddings ($E_{\text{WT}}$) were utilized.

##### 2.2 Dynamics Feature Extraction Using Anisotropic Network Models

Intrinsic protein dynamics were characterized using the Anisotropic Network Model (ANM) implemented in ProDy (v2.4)(11-13). Protein structures were represented as elastic networks in which Cα atoms within a cutoff distance were connected by uniform harmonic springs. The overall elastic potential of the network is defined as:

$$\begin{aligned} V=\frac{\gamma}{2}\sum_{i<j} \Gamma_{ij}\left( R_{ij}-R_{ij}^{0} \right)^{2}\#\left（ 2 \right） \end{aligned}$$

where $R_{ij}$ denotes the instantaneous distance between residues $i$ and $j$, $R_{ij}^{0}$ represents the equilibrium distance derived from the AlphaFold-predicted structure, and $\gamma$ is the uniform spring constant. The connectivity matrix $\Gamma_{ij}$ is a binary step function based on a spatial cutoff distance:

$$\Gamma_{ij}=\{\begin{matrix} 1, & R_{ij}^{0}<r_{c} \\ 0, & R_{ij}^{0}\geq r_{c} \end{matrix} (3)$$

where the cutoff radius $r_{c}$ was set to $15.0 A˚$

The Hessian matrix $H\in\mathbb{R}^{3N\times3N}$ was constructed from the second partial derivatives of the elastic potential. For interacting residue pairs, the off-diagonal super-elements of the Hessian were computed as:

$$\begin{aligned} H_{ij}=-\gamma\frac{r_{ij}r_{ij}^{T}}{\|r_{ij}{\|}^{2}}\# \left( 4 \right) \end{aligned}$$

where$r_{ij}$ denotes the displacement vector between residues $i$ and $j$.

Eigenvalue decomposition of the Hessian matrix ($HU = U\Lambda$) yielded the eigenvector matrix $U$, representing normal modes of motion, and the diagonal eigenvalue matrix $\Lambda$, corresponding to the squared frequencies of the modes.

After excluding the first six trivial modes corresponding to rigid-body translational and rotational motions, the first 30 nontrivial low-frequency modes and the last 30 high-frequency modes were retained. For each selected mode, the three Cartesian components corresponding to each residue were extracted and concatenated to generate residue-level dynamics embeddings. The low- and high-frequency embeddings were used as complementary node features for downstream graph learning.

##### 2.3 Adaptive Dynamics Fusion

To integrate complementary dynamics information from distinct ANMs, DynGeo-Pheno adopted an adaptive gating mechanism that selectively weighted multiple dynamics representations before graph-based learning. This design was motivated by the observation that low-frequency and high-frequency dynamics scales contributed unequally to residue-level pathogenicity across structural microenvironments.

Let $X\in\mathbb{R}^{N\times d_{seq}}$ denote sequence semantic embeddings from ESM-2, where $N$ is the number of residues and $d_{seq}$ is the sequence feature dimension. Two sets of residue-level dynamics embeddings derived from the low- and high-frequency ANM modes are denoted as $H_{low}\in\mathbb{R}^{N\times d_{dyn}}$ and $H_{high}\in\mathbb{R}^{N\times d_{dyn}}$. The two dynamics embeddings were first projected into a shared latent space,

$$\begin{matrix} {\overset{^}{H}}_{low} & =ReLU(H_{low}W_{f}+b_{f}) (5) \\ {\overset{^}{H}}_{high} & =ReLU(H_{high}W_{l}+b_{l}) (6) \end{matrix}$$

where $W_{f},W_{l}\in\mathbb{R}^{d_{dyn}\times d}$ are trainable projection matrices, $b_{f},b_{l}\in\mathbb{R}^{d}$ are bias vectors, and $d$ is the latent feature dimension.

The projected features were concatenated to compute gating coefficients,

$$G=\sigma\left( \left[ {\overset{^}{H}}_{low}\parallel{\overset{^}{H}}_{high} \right]W_{g}+b_{g} \right) (7)$$

Where $W_{g}\in\mathbb{R}^{2d\times d}$ is a learnable weight matrix, $b_{g}\in\mathbb{R}^{d}$ is a bias term, $\left[ \cdot\parallel\cdot\right]$ denotes feature-wise concatenation, and $\sigma(\cdot)$ is the sigmoid activation function.

The resulting gating coefficients $G\in\left[ 0,1 \right]^{N\times d}$ adaptively fused the two dynamics embeddings:

$$H_{fused}=G\odot{\overset{^}{H}}_{low}+\left( 1-G \right)\odot{\overset{^}{H}}_{high} (8)$$

where $\odot$ denotes element-wise multiplication. This formulation enabled residue specific weighting of low-frequency and high-frequency dynamics embeddings. Larger gating values retained information from the low-frequency representation ${\overset{^}{H}}_{low}$, while smaller values favored the high-frequency representation ${\overset{^}{H}}_{high}$.

Finally, the final node representation was constructed by concatenating the sequence and fused dynamics embeddings for downstream graph learning:

$$\begin{aligned} H_{node}=\left[ X\parallel H_{fused} \right]\#\left( 9 \right) \end{aligned}$$

where $H_{node}\in\mathbb{R}^{N\times\left( d_{seq}+d \right)}$. This hierarchical integration preserves full evolutionary information encoded by ESM-2 while introducing adaptively weighted dynamics priors, allowing the subsequent geometrical learning to jointly leverage sequence semantics and intrinsic protein dynamics for pathogenicity prediction.

#### 3. Geometric Neural Network Architecture: DynGeo-Pheno

##### 3.1 Graph Construction

Each protein structure was represented as an undirected spatial graph $G=\left( V,E \right)$, where the node set $V$ corresponds to individual amino acid residues centered on their C$\alpha$ atoms. The initial node feature matrix $H^{\left( 0 \right)}$, was obtained by concatenating the sequence embeddings and fused dynamics embeddings described in Section 2. To define the topological connectivity $E$, edges were established between residue pairs whose C$\alpha$ atoms were separated by a Euclidean distance of less than $10 A˚$ in three-dimensional space. For residue-level prediction, a local subgraph centered on each target residue was extracted using the *k*-nearest neighbor (kNN) algorithm with *k*=48. The resulting subgraphs were used as inputs to the graph neural network.

##### 3.2 Hybrid Residual GCN-GAT Encoder

The core encoder of DynGeo-Pheno employs a parallel architecture consisting of a modified Graph Convolutional Network (GCN) branch and a two-stage Graph Attention Network (GAT) branch.

For the GCN branch, a local residual feature propagation scheme is applied. Let $H^{\left( l \right)}$ denote the node representation matrix at layer $l$, and $P$ be the input adjacency matrix. The support representation $S^{\left( l \right)}$ is computed as:

$$\begin{aligned} S^{\left( l \right)}=\left( 1-\alpha\right)PH^{\left( l \right)}+\alpha H^{\left( l \right)}\#\left( 10 \right) \end{aligned}$$

where $\alpha$ is a fixed residual coefficient. The layer-wise forward propagation is then defined as:

$$\begin{aligned} H_{GCN}^{\left( l+1 \right)}=\sigma\left( \theta S^{\left( l \right)W^{\left( l \right)}}+\left( 1-\theta\right)S^{\left( l \right)W_{res}^{\left( l \right)}} \right)\#\left( 11 \right) \end{aligned}$$

where $W^{\left( l \right)}$ is the standard trainable convolution weight matrix, $W_{res}^{\left( l \right)}$ is a learnable residual projection matrix, and $\sigma\left( \cdot\right)$ denotes the ReLU activation function. The weighting coefficient $\theta$ is defined as $\theta=\min\left( 1.0,\log\left( \lambda+1.0 \right) \right)$, where $\lambda$ is a fixed hyperparameter.

Parallel to the GCN, the GAT branch employs a multi-head attention mechanism. In the first stage with $K$ independent attention heads, the attention coefficient $\alpha_{ij}^{k,\left( l \right)}$ between target residue $i$ and neighbor $j$ for head $k$ is calculated as:

$$\alpha_{ij}^{k,(l)}=\frac{exp(LeakyReLU(a_{k}^{(l)T}[W_{k}^{(l)}h_{i}^{(l)}\parallel W_{k}^{(l)}h_{j}^{(l)}]))}{\underset{u\in N_{i}}{\sum}exp(LeakyReLU(a_{k}^{(l)T}[W_{k}^{(l)}h_{i}^{(l)}\parallel W_{k}^{(l)}h_{u}^{(l)}]))} (12)$$

where $W_{k}^{\left( l \right)}$ is the projection matrix, $a_{k}^{\left( l \right)}$ is the attention weight vector, $\parallel$ denotes concatenation, and $\mathcal{N}_{\mathcal{i}}$ is the neighborhood of node $i$. The intermediate representation is formed by concatenating the outputs of the $K$ heads:

$$H_{multi}^{(l)}={\|}_{k=1}^{K}\sigma_{elu}(\underset{j\in N_{i}}{\sum}\alpha_{ij}^{k,(l)}W_{k}^{(l)}h_{j}^{(l)}) (13)$$

A second-stage attention layer then processes $H_{multi}^{\left( l \right)}$ using an analogous attention mechanism, but applies averaging instead of feature concatenation, to yield the final GAT representation $H_{GAT}^{\left( l+1 \right)}$.

##### 3.3 Parallel Feature Fusion and Prediction

Within each structural learning block, the heterogeneous representations generated by the GCN and GAT branches are dynamicly fused. Specifically, the outputs are concatenated and linearly projected to a unified embedding space. To facilitate seamless gradient backpropagation and preserve low-level physicochemical constraints, a block-level residual connection is introduced:

$$\begin{aligned} H^{\left( l+1 \right)}=\left[ H_{GAT}^{\left( l+1 \right)}\parallel H_{GCN}^{\left( l+1 \right)} \right]W_{fuse}^{\left( l \right)}+\text{Residual}^{\left( l \right)}\#\left( 14 \right) \end{aligned}$$

where $W_{fuse}^{\left( l \right)}$ is a linear transformation matrix regulating the fusion of topological and attentional features. To strictly align the feature dimensionalities, the skip connection term $\text{Residual}^{\left( l \right)}$ is adaptively defined as:

$${Residual}^{\left( l \right)}=\{\begin{matrix} H^{\left( l \right)}, & if d_{in}=d_{out} \\ H^{\left( l \right)}W_{skip}^{\left( l \right)}, & if d_{in}\neq d_{out} \end{matrix} (15)$$

where $W_{skip}^{\left( l \right)}$ is a learned projection matrix exclusively applied when the input dimension $d_{in}$ and output dimension $d_{out}$ differ.

The fully integrated node representations derived from the final layer are subsequently processed by a multi-layer perceptron (MLP) classifier with dropout regularization. The ultimate output layer employs a sigmoid activation function to yield $\hat{y}\in\left( 0,1 \right)$, quantifying the probability of a residue being disease-associated or structurally pathogenic.

#### 4. Implementation and Training Details

##### 4.1 Model Training

All models were implemented in Python 3.8.13 using the TensorFlow 2 framework and trained on a single NVIDIA GeForce RTX 2080 GPU. Model parameters were optimized using the RMSprop optimizer with an initial learning rate of ($1\times{10}^{-4}$). To mitigate overfitting, dropout regularization was applied with a dropout rate of 0.3 in graph neural network layers and 0.1 in the final multilayer perceptron classifier. Model development and hyperparameter optimization were performed using 10-fold stratified cross-validation on the training set. An independent model was trained for each fold using identical hyperparameter settings. Final predictions were obtained by majority voting across the ten models.

##### 4.2 Performance Evaluation

Model performance was evaluated using Precision, Recall, F1 score, Accuracy, Matthews correlation coefficient (MCC), the area under the receiver operating characteristic curve (AUROC), and the area under the precision–recall curve (AUPRC). Let TP, TN, FP, and FN denote the numbers of true positives, true negatives, false positives, and false negatives, respectively.

Precision, Recall, F1 score, and Accuracy were calculated as

$$\begin{aligned} \text{Precision}=\frac{\text{TP}}{\text{TP}+\text{FP}}\#\left( 16 \right) \end{aligned}$$

$$\begin{aligned} \text{Recall}=\frac{\text{TP}}{\text{TP}+\text{FN}}\#\left( 17 \right) \end{aligned}$$

$$\begin{aligned} \text{F1 Score}=2\times\frac{\text{Precision}\times\text{Recall}}{\text{Precision}+\text{Recall}}\#\left( 18 \right) \end{aligned}$$

$$\begin{aligned} \text{Accuracy}=\frac{\text{TP}+\text{TN}}{\text{TP}+\text{TN}+\text{FP}+\text{FN}}\#\left( 19 \right) \end{aligned}$$

Receiver operating characteristic (ROC) curves were generated by plotting the true positive rate (TPR) against the false positive rate (FPR),

$$\begin{aligned} \text{TPR}=\frac{\text{TP}}{\text{TP}+\text{FN}}\#\left( 20 \right) \end{aligned}$$

$$\begin{aligned} \text{FPR}=\frac{\text{FP}}{\text{FP}+\text{TN}}\#\left( 21 \right) \end{aligned}$$

and AUROC was calculated accordingly. Precision–recall (PR) curves were generated over the full range of decision thresholds, and the corresponding AUPRC was computed.

The Matthews correlation coefficient (MCC) was calculated as

$$\begin{aligned} MCC=\frac{TP\times TN-FP\times FN}{\sqrt{\left( TP+FP \right)\left( TP+FN \right)\left( TN+FP \right)\left( TN+FN \right)}}\#\left( 22 \right) \end{aligned}$$

#### 5. Biological and Dynamics Interpretation Analyses

All biological interpretation analyses were performed exclusively after model training and independent evaluation. Functional annotations, ENM-derived dynamics descriptors, enrichment statistics, and residue–residue cross correlation analyses were used solely for post hoc mechanistic interpretation and were never incorporated into model training or hyperparameter optimization. Consequently, no information leakage occurred between predictive modeling and biological interpretation.

##### 5.1 Biological Interpretation Analyses

To evaluate whether residues predicted as disease-associated preferentially localize within biologically meaningful regions, experimentally validated ligand-binding pockets, protein–protein interaction (PPI) interfaces and predicted cavities were mapped onto the structural datasets.

###### 5.1.1 Ligand-Binding Pocket Annotation (BioLiP)

Experimentally validated ligand-binding residues were obtained from the BioLiP database(14). After structural mapping and quality-control procedures, 11,905 ligand-binding pockets were retained. Following intersection with the benchmark datasets, 974 annotated pockets were mapped to proteins in the phosphorylation dataset (Table S6), and 1,000 pockets were mapped to proteins in the missense mutation dataset (Table S7).

###### 5.1.2 Protein-Protein Interaction Interface Annotation

Protein–protein interaction (PPI) annotations were derived from experimentally determined multimeric crystal structures of multimers in the RCSB Protein Data Bank (PDB)(15, 16). The curated interface annotations were mapped onto AlphaFold protein structures and intersected with the benchmark datasets. A total of 462 phosphorylation sites located within PPI interfaces were identified across 210 proteins (Table S8). For the missense mutation dataset, 412 interface-associated mutations from 128 proteins were retained after structural mapping and quality control (Table S9).

###### 5.1.3 Geometric Pocket Detection (Fpocket)

Protein surface cavities were identified using Fpocket(17). Cavity annotations were generated from protein geometry based on Voronoi tessellation and alpha-sphere clustering and subsequently mapped onto the benchmark datasets. The resulting cavity annotations are provided in Tables S10 and S11 for the phosphorylation and missense mutation datasets, respectively.

##### 5.2 Dynamics Interpretation Analyses

###### 5.2.1 ENM-Derived Dynamics Descriptors

Four residue-level ENM-derived descriptors, including root mean square fluctuation (RMSF), perturbation response scanning (PRS) effectiveness(18), mechanical stiffness(19), and residue cross-correlation (CC), were calculated using ProDy(12, 13).

The RMSF for residue $i$ is calculated as:

$$\begin{aligned} RMSF_{i}=\sqrt{\left\langle\Delta R_{i}^{2} \right\rangle}\#\left( 23 \right) \end{aligned}$$

where $\left\langle\Delta R_{i}^{2} \right\rangle$ denotes the mean-square fluctuation predicted from ENM normal modes.

The PRS effectiveness is computed as:

$$\begin{aligned} PRS_{i}=\frac{1}{N-1}\sum_{j\neq i} \left| \Delta R_{j}^{\left( i \right)} \right|\#\left( 24 \right) \end{aligned}$$

where $\Delta R_{j}^{\left( i \right)}$ represents the structural response of residue $j$ following a virtual perturbation applied to residue $i$.

The mean mechanical stiffness is defined as:

$$\begin{aligned} K_{i}=\frac{1}{N-1}\sum_{j\neq i} k_{ij}\#\left( 25 \right) \end{aligned}$$

where $k_{ij}$ denotes the effective spring constant between residues $i$ and $j$.

Residue-level cross-correlation (CC) was calculated from ENM normal modes using the ProDy function calcCrossCorr. For residue $i$, the CC score is defined as:

$$\begin{aligned} CC_{i}=\frac{1}{N-1}\sum_{j\neq i} \left| C_{ij} \right|\#\left( 26 \right) \end{aligned}$$

where $C_{ij}$ denotes the normalized cross-correlation between residues $i$ and $j$. Higher absolute CC values indicate stronger dynamics coupling. Distributions of RMSF, PRS effectiveness, mean stiffness, and CC were systematically compared between disease-associated and neutral residues in both benchmark datasets (Tables S12 and S13).

###### 5.2.2 Identification of Dynamics Hotspots

Dynamics hotspots were identified based on four residue-level ENM-derived descriptors: RMSF, CC, PRS effectiveness, and mechanical stiffness. For each protein, residues were independently ranked according to individual dynamics descriptors, and the top 20% of ranked residues were defined as dynamics hotspots. Specifically, residues exhibiting the lowest RMSF values were identified as rigidity hotspots, while residues with the highest mean absolute cross-correlation, PRS effectiveness, and mechanical stiffness scores were designated as communication, perturbation, and mechanical hotspots, respectively. Enrichment was evaluated using odds ratios (ORs) and density ratios (DRs).

###### 5.2.3 Dynamics Cross-Correlation Analyses

Pairwise residue cross-correlations were compared among disease-associated–disease-associated, neutral–neutral, and mixed perturbation pairs. To evaluate perturbation coupling with functional regions, perturbation sites overlapping BioLiP ligand-binding sites, Fpocket-predicted cavities, or PPI interfaces were excluded. For each remaining perturbation site, the mean absolute cross-correlation with all residues within the annotated functional region was calculated. Coupling strengths were then compared between disease-associated and neutral perturbation sites.

##### 5.3 Functional Enrichment Quantification

To quantify the extent to which disease-associated perturbations preferentially occupy functional regions, enrichment analyses were performed using both odds ratios (OR) and density ratios (DR). The odds ratio was defined as

$$\begin{aligned} OR=\frac{a/b}{c/d}\#\left( 27 \right) \end{aligned}$$

where $a$ and $b$ denote the numbers of disease-associated residues located inside and outside a functional region, respectively, whereas $c$and $d$represent the corresponding counts for background residues. To account for heterogeneity in annotation coverage among proteins, density ratios were additionally calculated as

$$\begin{aligned} DR=\frac{\rho_{region}}{\rho_{protein}}\#\left( 28 \right) \end{aligned}$$

where

$$\begin{aligned} \rho_{region}=\frac{N_{disease,region}}{N_{region}}\#\left( 29 \right) \end{aligned}$$

and

$$\begin{aligned} \rho_{protein}=\frac{N_{disease,protein}}{N_{protein}}\#\left( 30 \right) \end{aligned}$$

represent the disease-associated residue density within annotated functional regions and within the corresponding proteins, respectively. Enrichment analyses were performed independently for BioLiP ligand-binding sites, Fpocket cavities, PPI interfaces, and each category of dynamics hotspots (RMSF, CC, PRS effectiveness, and stiffness).

Because hotspot sizes differ among proteins, enrichment was additionally quantified using density ratios (DRs). Positive density ratio (Positive DR) measures the density of disease-associated residues within an annotated region relative to the average density across the corresponding protein, whereas negative density ratio (Negative DR) quantifies the same relationship for neutral residues.

### Table S3-S5

Table S3. Hyperparameter sensitivity analysis in DynGeo-Pheno. Performance was evaluated using AUROC under alternative configurations of ENM-derived dynamics representations, ESM embedding dimensionality, local structural neighborhood size, and AlphaFold confidence filtering. Bold values indicate the configuration adopted in the final model.

| Parameter category | Setting | Phos AUROC | Mut AUROC |
| --- | --- | --- | --- |
| Dynamics modes | 10 modes | 0.8227 | 0.8598 |
|  | 20 modes | 0.8353 | 0.8728 |
|  | **30 modes** | **0.8639** | **0.8968** |
|  | 40 modes | 0.8515 | 0.8178 |
| ESM dimension | 64 | 0.7688 | 0.8459 |
|  | **128** | **0.8529** | **0.9138** |
|  | 320 | 0.8184 | 0.8624 |
|  | 640 | 0.7981 | 0.8898 |
| KNN size | k = 12 | 0.7268 | 0.7996 |
|  | k = 24 | 0.8193 | 0.8603 |
|  | k = 36 | 0.8302 | 0.913 |
|  | **k = 48** | **0.8729** | **0.9261** |
| pLDDT filtering | 60–100 | 0.8313 | 0.8842 |
|  | **0–100** | **0.8679** | **0.9186** |

Table S4. Predictive performance and ensemble validation of DynGeo-Pheno on the prediction of disease associated phosphosite and missense mutation tasks. The table presents a detailed comparison of evaluation metrics among the 10-fold cross-validation average, the highest-performing single model from the cross-validation (fold 5 for phosphosite and fold 8 for mutation), and the final ensemble model. Performance is quantified using Accuracy, Precision, F1 Score, Recall, AUROC, and AUPRC. The ensemble model demonstrates improved stability and predictive power, particularly in the threshold-independent metrics (AUROC and AUPRC).

| Predictive Task | Model Configuration | ACC | Precision | F1 Score | Recall | AUROC | AUPRC |
| --- | --- | --- | --- | --- | --- | --- | --- |
| Phosphorylation | 10-Fold Mean | 0.7707 | 0.6123 | 0.6908 | 0.7965 | 0.8429 | 0.7205 |
|  | Best Single Model | 0.7941 | 0.6551 | 0.7232 | 0.8072 | 0.8587 | 0.7397 |
|  | **Ensemble Model** | **0.8039** | **0.6921** | 0.7161 | 0.7418 | **0.8729** | **0.7719** |
| Missense Mutation | 10-Fold Mean | 0.8506 | 0.7758 | 0.7891 | 0.8147 | 0.9223 | 0.8709 |
|  | Best Single Model | 0.8424 | 0.7204 | 0.794 | **0.8845** | 0.9266 | 0.8788 |
|  | **Ensemble Model** | **0.8811** | **0.8198** | **0.8288** | 0.8379 | **0.9368** | **0.8965** |

**Table S5. Ablation analysis of evolutionary and dynamics representations in DynGeo-Pheno.** Predictive performance was evaluated for disease-associated phosphosite prediction and pathogenic missense mutation prediction under different feature configurations. Performance was assessed using accuracy, precision, recall, F1 score, AUROC, and AUPRC. This table provides the complete quantitative results underlying the ablation analysis shown in Fig. 3D. The results show that sequence-derived ESM representations provide strong predictive performance, while explicit incorporation of dynamics yields complementary effects that are reflected not only in AUROC but also in AUPRC across the two prediction tasks.

| Predictive Task | Category | ACC | Precision | F1_score | Recall | AUROC | AUPRC |
| --- | --- | --- | --- | --- | --- | --- | --- |
| Phosphorylation | **Full Model** | **0.7707** | 0.6123 | **0.6908** | **0.7965** | **0.8429** | **0.7205** |
|  | ESM + High | 0.7561 | 0.6062 | 0.6805 | 0.7791 | 0.8311 | 0.7054 |
|  | ESM +Low | 0.7520 | 0.5998 | 0.6779 | 0.7833 | 0.8293 | 0.7036 |
|  | ESM only | 0.7686 | 0.6317 | 0.6861 | 0.7588 | 0.8360 | 0.7110 |
|  | ANM Low + High | 0.6638 | 0.4915 | 0.5944 | 0.7601 | 0.7293 | 0.5401 |
|  | ANM Low only | 0.6471 | 0.4847 | 0.5954 | 0.7768 | 0.7289 | 0.5446 |
|  | ANM High only | 0.5595 | 0.4180 | 0.5345 | 0.7575 | 0.6587 | 0.4713 |
| Missense Mutation | **Full Model** | 0.8506 | 0.7758 | **0.7891** | 0.8147 | **0.9223** | **0.8709** |
|  | ESM + High | 0.8566 | 0.8178 | 0.7821 | 0.7494 | 0.9131 | 0.8504 |
|  | ESM +Low | 0.8509 | 0.7718 | 0.7873 | 0.8034 | 0.9138 | 0.8545 |
|  | ESM only | 0.8610 | 0.8281 | 0.7877 | 0.7511 | 0.9184 | 0.8658 |
|  | ANM Low + High | 0.6087 | 0.4643 | 0.6136 | 0.9047 | 0.7928 | 0.6683 |
|  | ANM Low only | 0.7663 | 0.6377 | 0.6850 | 0.7400 | 0.8449 | 0.7756 |
|  | ANM High only | 0.6834 | 0.5287 | 0.6095 | 0.7195 | 0.7474 | 0.6182 |

### Legends for Tables S1-2, S6-13

Table S1. Benchmark dataset for disease-associated phosphosite prediction. Columns include the UniProt accession (ACC_ID), source database (Database), phosphorylated residue and position (Phosphosite), binary class label (True_Label; 1 = disease-associated, 0 = neutral), and AlphaFold residue confidence (AF_pLDDT). This dataset was used for model training, validation, and independent testing.

Table S2. Benchmark dataset for pathogenic missense mutation prediction. Columns include the UniProt accession (ACC_ID), residue position (Position), wild-type amino acid (Wild_AA), mutant amino acid (Mut_AA). This dataset was used for model training, validation, and independent testing.

Table S6. BioLiP ligand-binding pocket annotations mapped to the phosphosite benchmark dataset. Each row corresponds to one experimentally validated ligand-binding pocket. In addition to the pocket identifier and constituent residues, the table reports phosphosites located inside each BioLiP pocket (In_Pocket_Positions) together with their DynGeo-Pheno prediction scores (In_Pocket_Scores), as well as phosphosites outside the pocket (Out_Pocket_Positions) and their corresponding prediction scores (Out_Pocket_Scores). These annotations were used for the structural interpretation analyses shown in Fig. 4A and Fig. 4B.

Table S7. BioLiP ligand-binding pocket annotations mapped to the missense mutation benchmark dataset. The table reports pathogenic missense mutations located inside (In_Pocket_Positions) and outside (Out_Pocket_Positions) of validated BioLiP ligand-binding pockets together with their corresponding DynGeo-Pheno prediction scores. These annotations were used for the structural interpretation analyses shown in Fig. 4A and Fig. 4B.

Table S8. PPI interface annotations for the phosphosite benchmark dataset. Each row corresponds to one phosphosite. Besides the common site information described in Table S1, the table includes the DynGeo-Pheno prediction score and a binary indicator (is_PPI_interface) specifying whether the residue is located within a PPI interface. These annotations were used for the analyses presented in Fig. 4A and Fig. 4B.

Table S9. PPI interface annotations for the missense mutation benchmark dataset. Each row corresponds to one missense mutation. In addition to the mutation information described in Table S2, the table reports the DynGeo-Pheno prediction score and whether the mutated residue is located within a PPI interface (is_PPI_interface). These annotations were used for the analyses presented in Fig. 4A and Fig. 4B.

Table S10. Fpocket geometric cavity annotations mapped to the phosphosite benchmark dataset. Each row corresponds to one predicted geometric cavity. The table reports phosphosites located inside (In_Pocket_Positions) and outside (Out_Pocket_Positions) of each predicted cavity together with their corresponding DynGeo-Pheno prediction scores. These annotations were used for the analyses presented in Fig. 4A and Fig. 4B.

Table S11. Fpocket geometric cavity annotations mapped to the missense mutation benchmark dataset. Each row corresponds to one predicted geometric cavity. The table reports pathogenic missense mutations located inside (In_Pocket_Positions) and outside (Out_Pocket_Positions) of each predicted cavity together with their corresponding DynGeo-Pheno prediction scores. These annotations were used for the analyses presented in Fig. 4A and Fig. 4B.

Table S12. ENM-derived dynamics descriptors for the phosphosite benchmark dataset. In addition to the site information reported in Table S1, the table provides residue-level dynamics descriptors derived from ANMs, including RMSF, cross-correlation (CC), perturbation response scanning (PRS) descriptors, and mechanical stiffness computed from the complete mode spectrum (_all), the lowest-frequency 30 ANM modes (_first30), or the highest-frequency 30 ANM modes (_last30). These descriptors were used for the dynamics analyses presented in Fig. 4C–F.

Table S13. ENM-derived dynamics descriptors for the missense mutation benchmark dataset. In addition to the mutation information reported in Table S2, the table provides the same set of ANM-derived dynamics descriptors described in Table S11. These descriptors were used for the dynamics analyses presented in Fig. 4C–F.

### Supplementary Figures

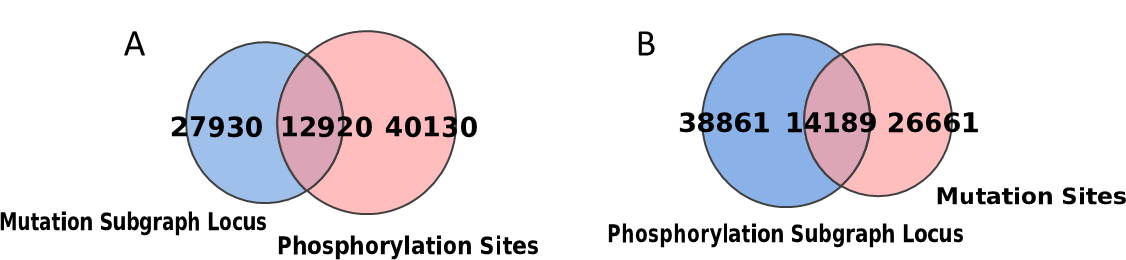

Figure S1. Structural overlap between phosphosite and mutation neighborhoods. (A) Venn diagram showing the overlap between mutation-centered KNN (k = 48) residue neighborhoods and phosphosite loci. Each mutation-centered subgraph was constructed by selecting the 48 nearest neighboring residues surrounding the mutation site in three-dimensional space. The overlap quantifies the extent to which phosphosites are contained within structural neighborhoods centered on mutation sites. (B) Reciprocal analysis showing the overlap between phosphosite-centered KNN (k = 48) residue neighborhoods and mutation loci. Each phosphosite-centered subgraph was generated using the same KNN strategy.

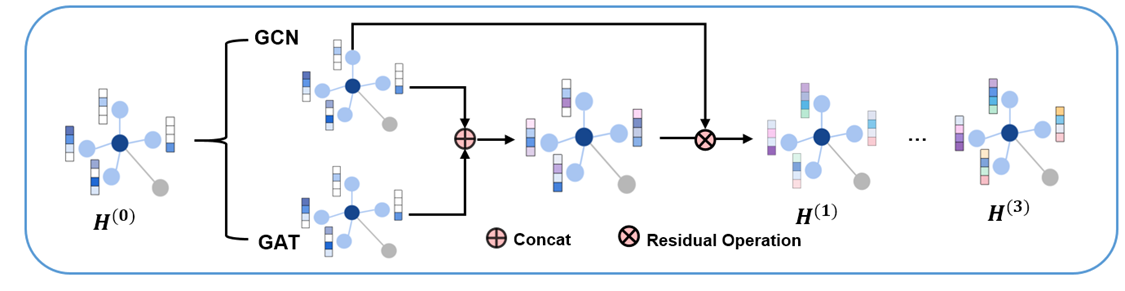

Figure S2. Architecture of each graph neural network block. At each layer, GCN and GAT features were extracted in parallel and concatenated, followed by residual connections to facilitate information propagation and stabilize optimization across multiple graph convolution layers.

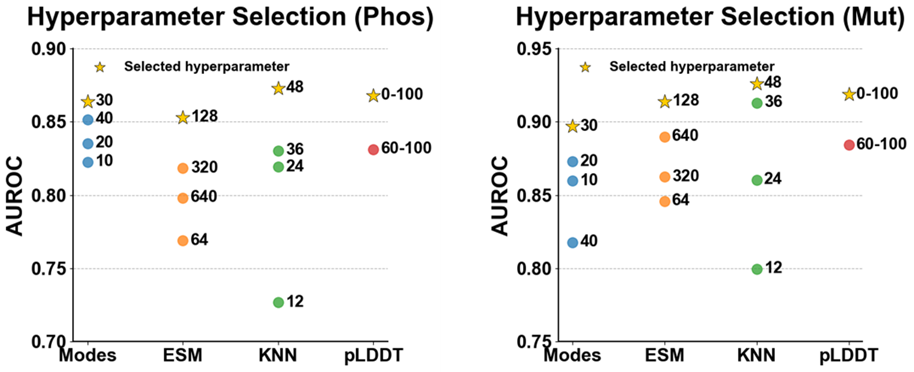

Figure S3. Hyperparameter optimization for DynGeo-Pheno. Hyperparameter selection was performed independently for phosphosite prediction (left) and missense mutation prediction (right). Four hyperparameters were evaluated, including the number of ANM modes, ESM embedding dimensionality after PCA, the number of K-nearest neighbors used for graph construction, and the pLDDT confidence threshold. Yellow stars indicate the final parameter combination adopted in the study (30 ANM modes, 128-dimensional ESM embeddings, K = 48, and no pLDDT filtering).

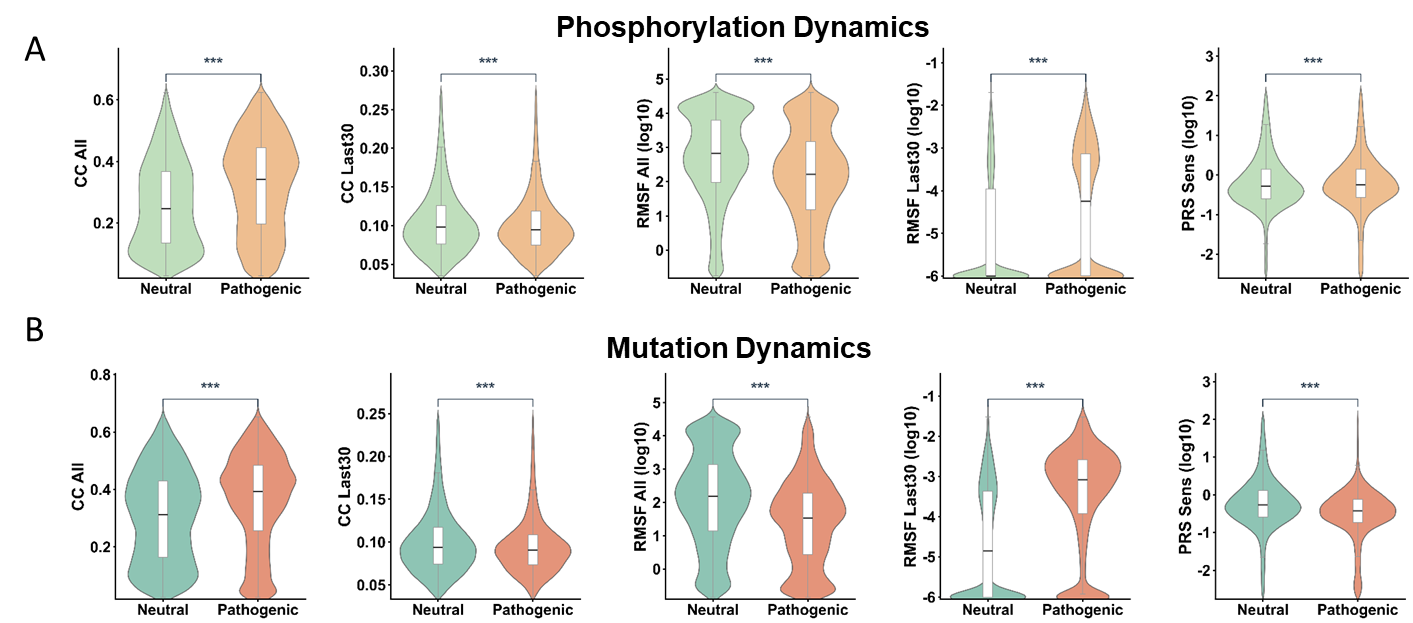

Figure S4. Additional comparisons of ENM-derived dynamics descriptors between disease-associated and neutral perturbations. Violin plots illustrate the distributions of additional dynamics descriptors that were not included in the main figures. Panel A correspond to phosphosites, whereas panels B correspond to pathogenic missense mutations. Comparisons include whole-spectrum and high-frequency cross-correlation (CC), whole-spectrum RMSF and high-frequency RMSF, and PRS sensitivity. Statistical significance was assessed using two-sided Mann–Whitney U tests, with significance levels indicated by asterisks. These complementary analyses further demonstrate that disease-associated residues exhibit characteristic dynamics signatures across multiple ENM-derived metrics beyond those presented in the main text.

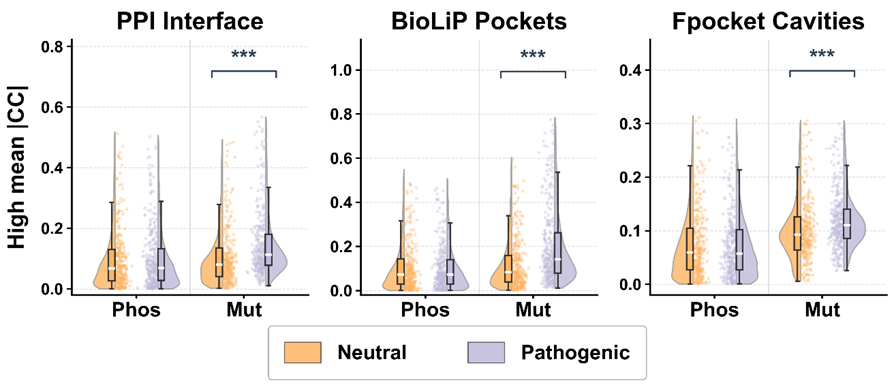

Figure S5. Dynamics coupling between disease-associated residues and functional regions under the highest-frequency 30 ANM modes. Disease-associated residues located outside PPI interfaces, BioLiP ligand-binding pockets, and Fpocket cavities exhibited significantly stronger dynamics coupling to these functional regions than neutral residues. Directly annotated functional residues were excluded prior to analysis, indicating that disease-associated perturbations remain functionally connected to canonical functional sites through long-range dynamics communication rather than simple spatial colocalization.

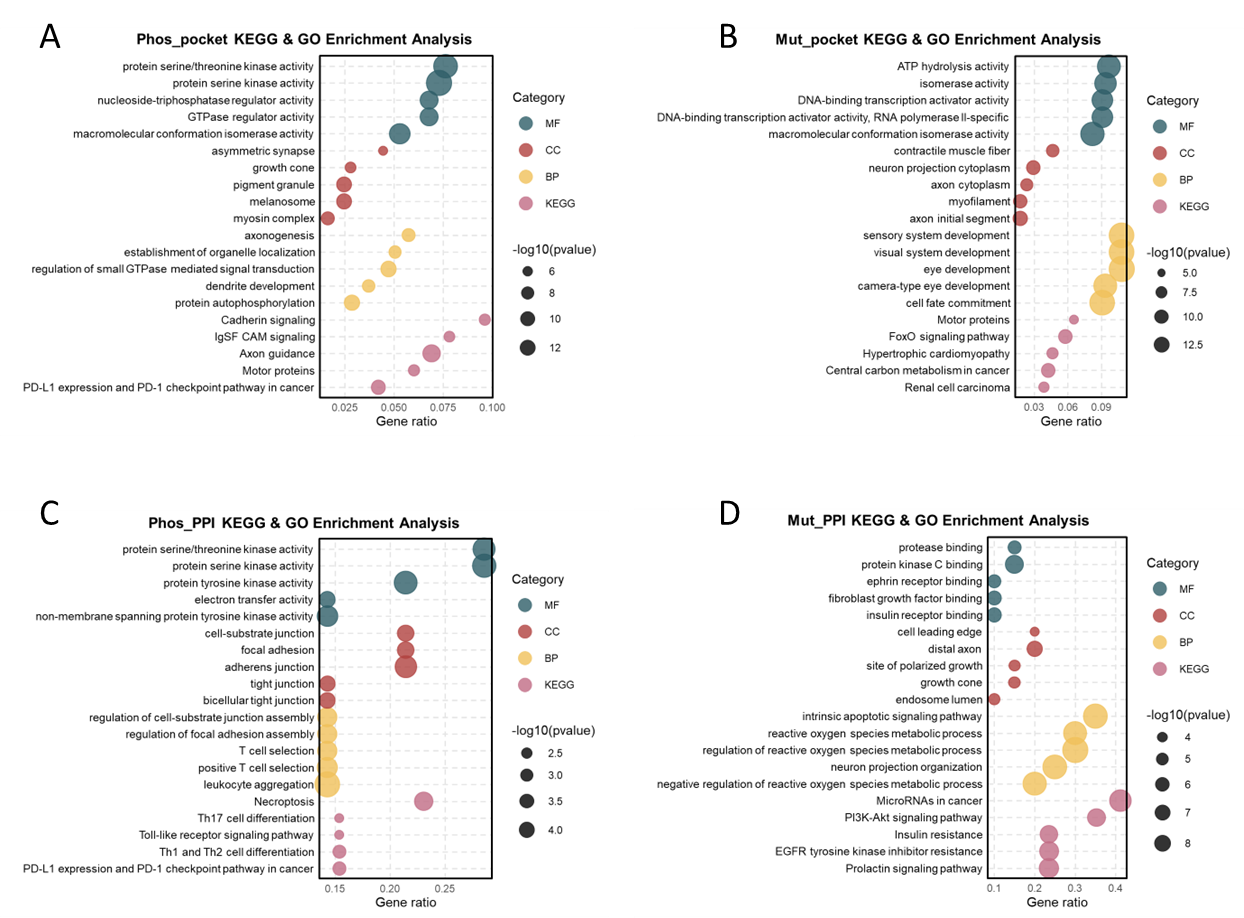

Figure S6. Functional enrichment analyses of proteins harboring structurally enriched disease-associated residues. (A) GO and KEGG enrichment of 852 proteins containing disease-associated phosphosites significantly enriched in ligand-binding pockets. (B) GO and KEGG enrichment of 639 proteins containing pathogenic mutations significantly enriched in ligand-binding pockets. (C) GO and KEGG enrichment of 15 proteins containing disease-associated phosphosites significantly enriched at PPI interfaces. (D) GO and KEGG enrichment of 20 proteins containing pathogenic mutations significantly enriched at PPI interfaces.

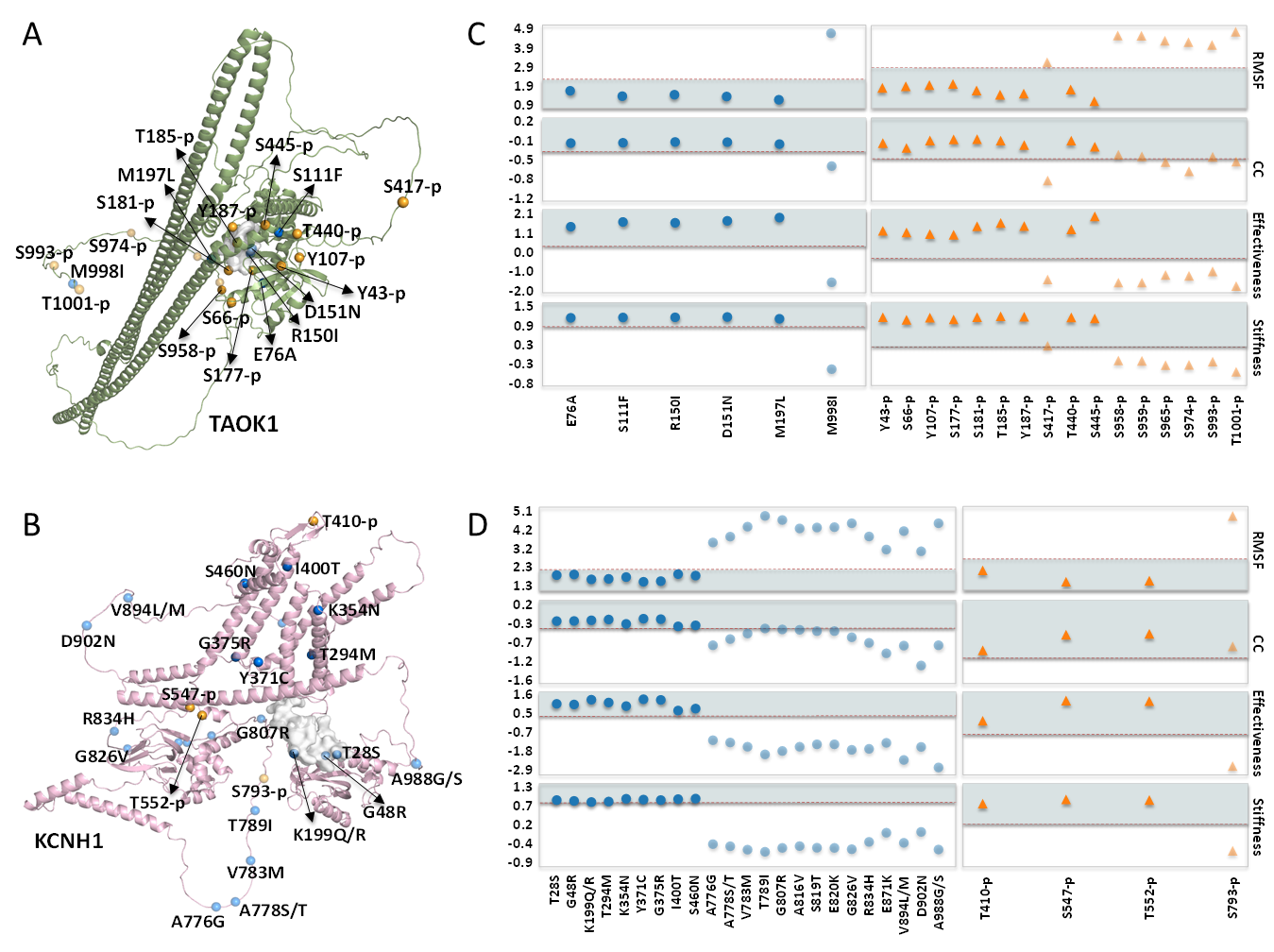

Figure S7. Supplementary case studies. (A, B) Structural representations and Fpocket-predicted pockets of the TAOK1 (A) and KCNH1 (B) systems. For both systems, only the single top-scoring Fpocket-predicted pocket overlapping with the positive sites is displayed. (C, D) Corresponding metric distribution plots for the TAOK1 (C) and KCNH1 (D) systems. All visualization, coloring, and transparency standards are consistent with those in Fig. 5.
