## Supporting Materials and Methods, Supporting Figures S1-S7, Supporting Tables S1-S13 for "Dynamics-aware geometric learning predicts disease-associated molecular perturbations": SUPPORTING_INFORMATION.pdf

For missense mutation prediction, mutation-induced sequence perturbations were represented by the difference between the PCA-reduced embeddings of the wild-type and mutant residues:

$$\Delta E = E_{\mathrm{WT}} - E_{\mathrm{Mut}} \quad (1)$$

where $E_{\mathrm{WT}}$ and $E_{\mathrm{Mut}} \in \mathbb{R}^{128}$ represent the PCA-reduced embeddings of the wild-type and mutant residues, respectively. For phosphorylation prediction, only wild-type embeddings ($E_{\mathrm{WT}}$) were utilized.

#### 2.2 Dynamics Feature Extraction Using Anisotropic Network Models

Intrinsic protein dynamics were characterized using the Anisotropic Network Model (ANM) implemented in ProDy (v2.4)(11-13). Protein structures were represented as elastic networks in which Cα atoms within a cutoff distance were connected by uniform harmonic springs. The overall elastic potential of the network is defined as:

$$V = \frac{\gamma}{2}\sum_{i<j}\Gamma_{ij}\left(R_{ij} - R_{ij}^{0}\right)^{2} \tag{2}$$

where $R_{ij}$ denotes the instantaneous distance between residues $i$ and $j$, $R_{ij}^{0}$ represents the equilibrium distance derived from the AlphaFold-predicted structure, and $\gamma$ is the uniform spring constant. The connectivity matrix $\Gamma_{ij}$ is a binary step function based on a spatial cutoff distance:

$$\Gamma_{ij} = \begin{cases} 1, & R_{ij}^{0} < r_c \\ 0, & R_{ij}^{0} \geq r_c \end{cases} \tag{3}$$

where the cutoff radius $r_c$ was set to 15.0 A°

The Hessian matrix $H \in \mathbb{R}^{3N\times 3N}$ was constructed from the second partial derivatives of the elastic potential. For interacting residue pairs, the off-diagonal super-elements of the Hessian were computed as:

$$H_{ij} = -\gamma\frac{r_{ij}r_{ij}^{T}}{\|r_{ij}\|^{2}} \tag{4}$$

where $r_{ij}$ denotes the displacement vector between residues $i$ and $j$.

Eigenvalue decomposition of the Hessian matrix ($HU = U\Lambda$) yielded the eigenvector matrix $U$, representing normal modes of motion, and the diagonal eigenvalue matrix $\Lambda$, corresponding to the squared frequencies of the modes.

After excluding the first six trivial modes corresponding to rigid-body translational and rotational motions, the first 30 nontrivial low-frequency modes and the last 30 high-frequency modes were retained. For each selected mode, the three Cartesian components corresponding to each residue were extracted and concatenated to generate residue-level dynamics embeddings. The low- and high-frequency embeddings were used as complementary node features for downstream graph learning.

#### 2.3 Adaptive Dynamics Fusion

To integrate complementary dynamics information from distinct ANMs, DynGeo-Pheno adopted an adaptive gating mechanism that selectively weighted multiple dynamics representations before graph-based learning. This design was motivated by the observation that low-frequency and high-frequency dynamics scales contributed unequally to residue-level pathogenicity across structural microenvironments.

Let $X \in \mathbb{R}^{N\times d_{seq}}$ denote sequence semantic embeddings from ESM-2, where $N$ is the number of residues and $d_{seq}$ is the sequence feature dimension. Two sets of residue-level dynamics embeddings derived from the low- and high-frequency ANM modes are denoted as $H_{low} \in \mathbb{R}^{N\times d_{dyn}}$ and $H_{high} \in \mathbb{R}^{N\times d_{dyn}}$. The two dynamics embeddings were first projected into a shared latent space,

$$\hat{H}_{low} = ReLU(H_{low}W_f + b_f) \tag{5}$$

$$\hat{H}_{high} = ReLU(H_{high}W_l + b_l) \tag{6}$$

where $W_f, W_l \in \mathbb{R}^{d_{dyn}\times d}$ are trainable projection matrices, $b_f, b_l \in \mathbb{R}^d$ are bias vectors, and $d$ is the latent feature dimension.

The projected features were concatenated to compute gating coefficients,

$$G = \sigma\left(\left[\hat{H}_{low} \parallel \hat{H}_{high}\right] W_g + b_g\right) \tag{7}$$

Where $W_g \in \mathbb{R}^{2d\times d}$ is a learnable weight matrix, $b_g \in \mathbb{R}^d$ is a bias term, $[\cdot\|\cdot]$ denotes feature-wise concatenation, and $\sigma(\cdot)$ is the sigmoid activation function.

The resulting gating coefficients $G \in [0,1]^{N\times d}$ adaptively fused the two dynamics embeddings:

$$H_{fused} = G \odot \hat{H}_{low} + (1 - G) \odot \hat{H}_{high} \tag{8}$$

where $\odot$ denotes element-wise multiplication. This formulation enabled residue specific weighting of low-frequency and high-frequency dynamics embeddings. Larger gating values retained information from the low-frequency representation $\hat{H}_{low}$, while smaller values favored the high-frequency representation $\hat{H}_{high}$.

Finally, the final node representation was constructed by concatenating the sequence and fused dynamics embeddings for downstream graph learning:

$$H_{node} = \left[X \parallel H_{fused}\right] \tag{9}$$

where $H_{node} \in \mathbb{R}^{N\times(d_{seq}+d)}$. This hierarchical integration preserves full evolutionary information encoded by ESM-2 while introducing adaptively weighted dynamics priors, allowing the subsequent geometrical learning to jointly leverage sequence semantics and intrinsic protein dynamics for pathogenicity prediction.

### 3. Geometric Neural Network Architecture: DynGeo-Pheno

#### 3.1 Graph Construction

Each protein structure was represented as an undirected spatial graph $G = (V, E)$, where the node set $V$ corresponds to individual amino acid residues centered on their Cα atoms. The initial node feature matrix $H^{(0)}$, was obtained by concatenating the sequence embeddings and fused dynamics embeddings described in Section 2. To define the topological connectivity $E$, edges were established between residue pairs whose Cα atoms were separated by a Euclidean distance of less than 10 A° in three-dimensional space. For residue-level prediction, a local subgraph centered on each target residue was extracted using the *k*-nearest neighbor (kNN) algorithm with *k*=48. The resulting subgraphs were used as inputs to the graph neural network.

#### 3.2 Hybrid Residual GCN-GAT Encoder

The core encoder of DynGeo-Pheno employs a parallel architecture consisting of a modified Graph Convolutional Network (GCN) branch and a two-stage Graph Attention Network (GAT) branch.

For the GCN branch, a local residual feature propagation scheme is applied. Let $H^{(l)}$ denote the node representation matrix at layer $l$, and $P$ be the input adjacency matrix. The support representation $S^{(l)}$ is computed as:

$$S^{(l)} = (1-\alpha)PH^{(l)} + \alpha H^{(l)} \tag{10}$$

where $\alpha$ is a fixed residual coefficient. The layer-wise forward propagation is then defined as:

$$H_{GCN}^{(l+1)} = \sigma\left(\theta S^{(l)}W^{(l)} + (1-\theta)S^{(l)}W_{res}^{(l)}\right) \tag{11}$$

where $W^{(l)}$ is the standard trainable convolution weight matrix, $W_{res}^{(l)}$ is a learnable residual projection matrix, and $\sigma(\cdot)$ denotes the ReLU activation function. The weighting coefficient $\theta$ is defined as $\theta = \min(1.0, \log(\lambda + 1.0))$, where $\lambda$ is a fixed hyperparameter.

Parallel to the GCN, the GAT branch employs a multi-head attention mechanism. In the first stage with $K$ independent attention heads, the attention coefficient $\alpha_{ij}^{k,(l)}$ between target residue $i$ and neighbor $j$ for head $k$ is calculated as:

$$\alpha_{ij}^{k,(l)} = \frac{exp(LeakyReLU(a_k^{(l)T}[W_k^{(l)}h_i^{(l)} \parallel W_k^{(l)}h_j^{(l)}]))}{\sum_{u \in N_i} exp(LeakyReLU(a_k^{(l)T}[W_k^{(l)}h_i^{(l)} \parallel W_k^{(l)}h_u^{(l)}]))} \tag{12}$$

where $W_k^{(l)}$ is the projection matrix, $a_k^{(l)}$ is the attention weight vector, $\parallel$ denotes concatenation, and $\mathcal{N}_i$ is the neighborhood of node $i$. The intermediate representation is formed by concatenating the outputs of the $K$ heads:

$$H_{multi}^{(l)} = \parallel_{k=1}^{K} \sigma_{elu}\left(\sum_{j \in N_i} \alpha_{ij}^{k,(l)} W_k^{(l)} h_j^{(l)}\right) \tag{13}$$

A second-stage attention layer then processes $H_{multi}^{(l)}$ using an analogous attention mechanism, but applies averaging instead of feature concatenation, to yield the final GAT representation $H_{GAT}^{(l+1)}$.

#### 3.3 Parallel Feature Fusion and Prediction

Within each structural learning block, the heterogeneous representations generated by the GCN and GAT branches are dynamicly fused. Specifically, the outputs are concatenated and linearly projected to a unified embedding space. To facilitate seamless gradient backpropagation and preserve low-level physicochemical constraints, a block-level residual connection is introduced:

$$H^{(l+1)} = \left[H_{GAT}^{(l+1)} \parallel H_{GCN}^{(l+1)}\right] W_{fuse}^{(l)} + \text{Residual}^{(l)} \tag{14}$$

where $W_{fuse}^{(l)}$ is a linear transformation matrix regulating the fusion of topological and attentional features. To strictly align the feature dimensionalities, the skip connection term $\text{Residual}^{(l)}$ is adaptively defined as:

$$Residual^{(l)} = \begin{cases} H^{(l)}, & if\ d_{in} = d_{out} \\ H^{(l)} W_{skip}^{(l)}, & if\ d_{in} \neq d_{out} \end{cases} \tag{15}$$

where $W_{skip}^{(l)}$ is a learned projection matrix exclusively applied when the input dimension $d_{in}$

and output dimension $d_{out}$ differ.

The fully integrated node representations derived from the final layer are subsequently processed by a multi-layer perceptron (MLP) classifier with dropout regularization. The ultimate output layer employs a sigmoid activation function to yield $\hat{y} \in (0,1)$, quantifying the probability of a residue being disease-associated or structurally pathogenic.

### 4. Implementation and Training Details

#### 4.1 Model Training

All models were implemented in Python 3.8.13 using the TensorFlow 2 framework and trained on a single NVIDIA GeForce RTX 2080 GPU. Model parameters were optimized using the RMSprop optimizer with an initial learning rate of $(1 \times 10^{-4})$. To mitigate overfitting, dropout regularization was applied with a dropout rate of 0.3 in graph neural network layers and 0.1 in the final multilayer perceptron classifier. Model development and hyperparameter optimization were performed using 10-fold stratified cross-validation on the training set. An independent model was trained for each fold using identical hyperparameter settings. Final predictions were obtained by majority voting across the ten models.

#### 4.2 Performance Evaluation

Model performance was evaluated using Precision, Recall, F1 score, Accuracy, Matthews correlation coefficient (MCC), the area under the receiver operating characteristic curve (AUROC), and the area under the precision–recall curve (AUPRC). Let TP, TN, FP, and FN denote the numbers of true positives, true negatives, false positives, and false negatives, respectively.

Precision, Recall, F1 score, and Accuracy were calculated as

$$\text{Precision} = \frac{\text{TP}}{\text{TP} + \text{FP}} \tag{16}$$

$$\text{Recall} = \frac{\text{TP}}{\text{TP} + \text{FN}} \tag{17}$$

$$\text{F1 Score} = 2 \times \frac{\text{Precision} \times \text{Recall}}{\text{Precision} + \text{Recall}} \tag{18}$$

$$\text{Accuracy} = \frac{\text{TP} + \text{TN}}{\text{TP} + \text{TN} + \text{FP} + \text{FN}} \tag{19}$$

Receiver operating characteristic (ROC) curves were generated by plotting the true positive rate (TPR) against the false positive rate (FPR),

$$\text{TPR} = \frac{\text{TP}}{\text{TP} + \text{FN}} \tag{20}$$

$$\text{FPR} = \frac{\text{FP}}{\text{FP} + \text{TN}} \tag{21}$$

and AUROC was calculated accordingly. Precision–recall (PR) curves were generated over the full range of decision thresholds, and the corresponding AUPRC was computed.

The RMSF for residue $i$ is calculated as:

$$RMSF_i = \sqrt{\langle \Delta R_i^2 \rangle} \tag{23}$$

where $\langle \Delta R_i^2 \rangle$ denotes the mean-square fluctuation predicted from ENM normal modes.

The PRS effectiveness is computed as:

$$PRS_i = \frac{1}{N-1} \sum_{j \neq i} \left| \Delta R_j^{(i)} \right| \tag{24}$$

where $\Delta R_j^{(i)}$ represents the structural response of residue $j$ following a virtual perturbation applied to residue $i$.

The mean mechanical stiffness is defined as:

$$DR = \frac{\rho_{region}}{\rho_{protein}} \tag{28}$$

where

$$\rho_{region} = \frac{N_{disease,region}}{N_{region}} \tag{29}$$

and

$$\rho_{protein} = \frac{N_{disease,protein}}{N_{protein}} \tag{30}$$

represent the disease-associated residue density within annotated functional regions and within the corresponding proteins, respectively. Enrichment analyses were performed independently for BioLiP ligand-binding sites, Fpocket cavities, PPI interfaces, and each category of dynamics hotspots (RMSF, CC, PRS effectiveness, and stiffness).

Table S13. ENM-derived dynamics descriptors for the missense mutation benchmark dataset. In addition to the mutation information reported in Table S2, the table provides the same set of ANM-derived dynamics descriptors described in Table S11. These descriptors were used for the dynamics analyses presented in Fig. 4C–F.

#### Supplementary Figures

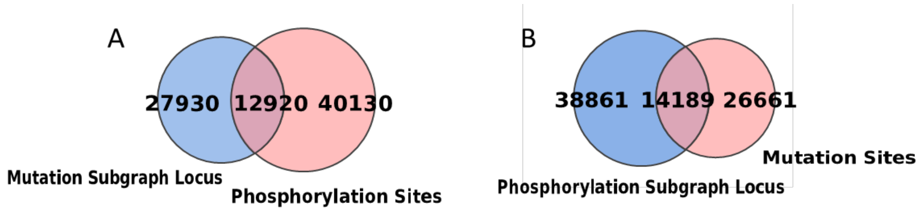


Figure S1. Structural overlap between phosphosite and mutation neighborhoods. (A) Venn diagram showing the overlap between mutation-centered KNN (k = 48) residue neighborhoods and phosphosite loci. Each mutation-centered subgraph was constructed by selecting the 48 nearest neighboring residues surrounding the mutation site in three-dimensional space. The overlap quantifies the extent to which phosphosites are contained within structural neighborhoods centered on mutation sites. (B) Reciprocal analysis showing the overlap between phosphosite-centered KNN (k = 48) residue neighborhoods and mutation loci. Each phosphosite-centered subgraph was generated using the same KNN strategy.

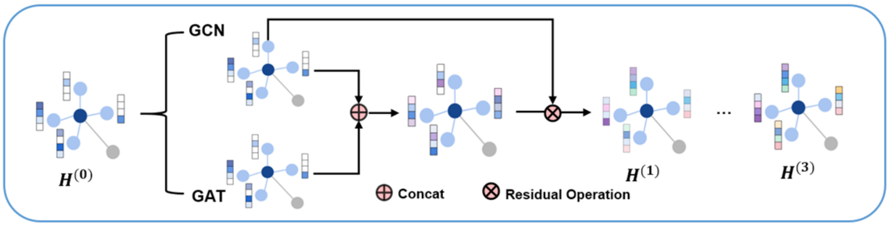


Figure S2. Architecture of each graph neural network block. At each layer, GCN and GAT features were extracted in parallel and concatenated, followed by residual connections to facilitate information propagation and stabilize optimization across multiple graph convolution layers.

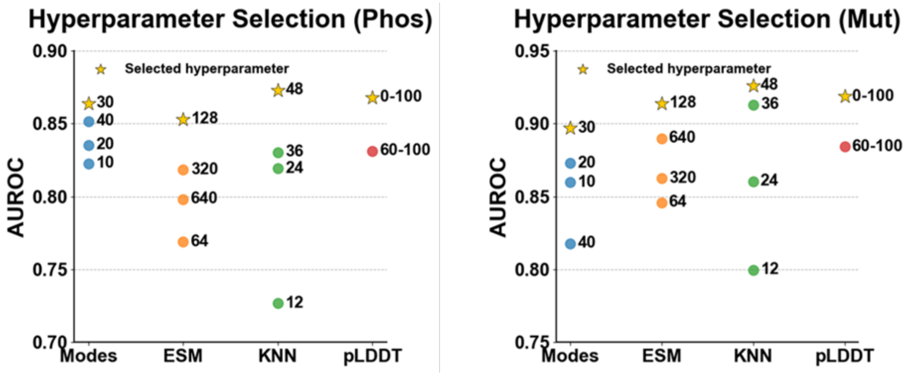


Figure S3. Hyperparameter optimization for DynGeo-Pheno. Hyperparameter selection was performed independently for phosphosite prediction (left) and missense mutation prediction (right). Four hyperparameters were evaluated, including the number of ANM modes, ESM embedding dimensionality after PCA, the number of K-nearest neighbors used for graph construction, and the pLDDT confidence threshold. Yellow stars indicate the final parameter combination adopted in the study (30 ANM modes, 128-dimensional ESM embeddings, K = 48, and no pLDDT filtering).

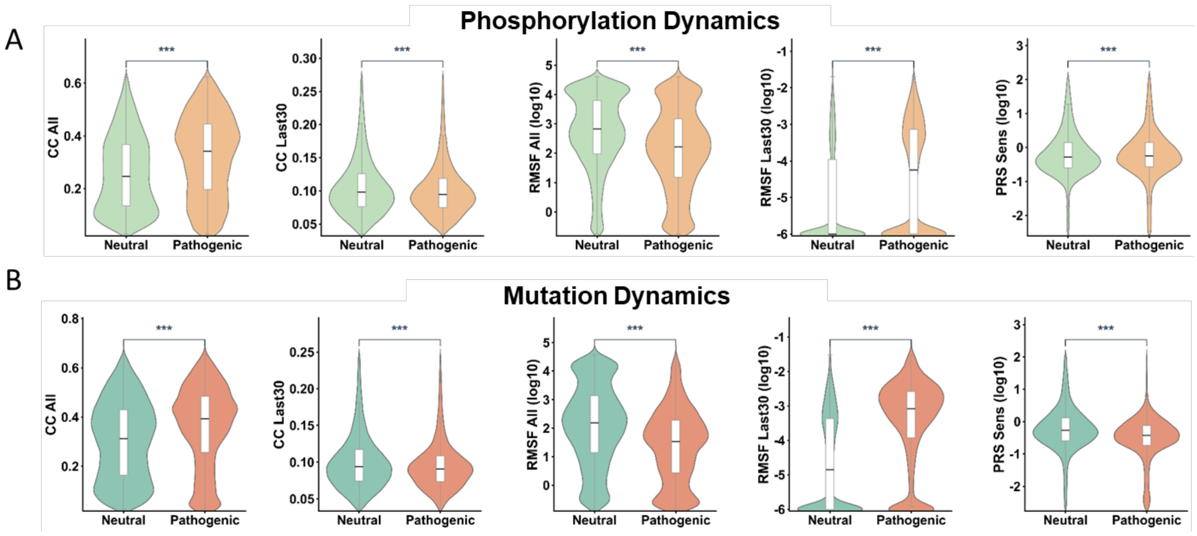


Figure S4. Additional comparisons of ENM-derived dynamics descriptors between disease-associated and neutral perturbations. Violin plots illustrate the distributions of additional dynamics descriptors that were not included in the main figures. Panel A correspond to phosphosites, whereas panels B correspond to pathogenic missense mutations. Comparisons include whole-spectrum and high-frequency cross-correlation (CC), whole-spectrum RMSF and high-frequency RMSF, and PRS sensitivity. Statistical significance was assessed using two-sided Mann–Whitney U tests, with significance levels indicated by asterisks. These complementary analyses further demonstrate that disease-associated residues exhibit characteristic dynamics signatures across multiple ENM-derived metrics beyond those presented in the main text.

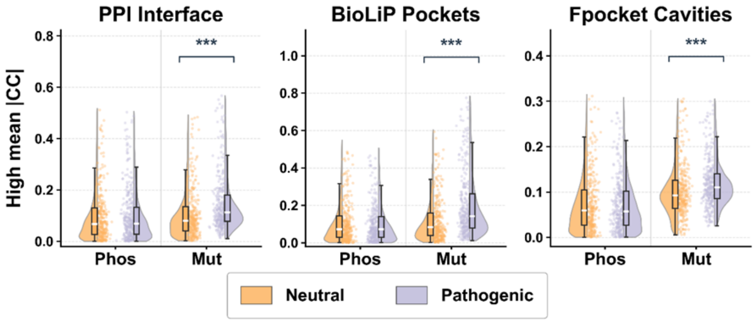


Figure S5. Dynamics coupling between disease-associated residues and functional regions under the highest-frequency 30 ANM modes. Disease-associated residues located outside PPI interfaces, BioLiP ligand-binding pockets, and Fpocket cavities exhibited significantly stronger dynamics coupling to these functional regions than neutral residues. Directly annotated functional residues were excluded prior to analysis, indicating that disease-associated perturbations remain functionally connected to canonical functional sites through long-range dynamics communication rather than simple spatial colocalization.

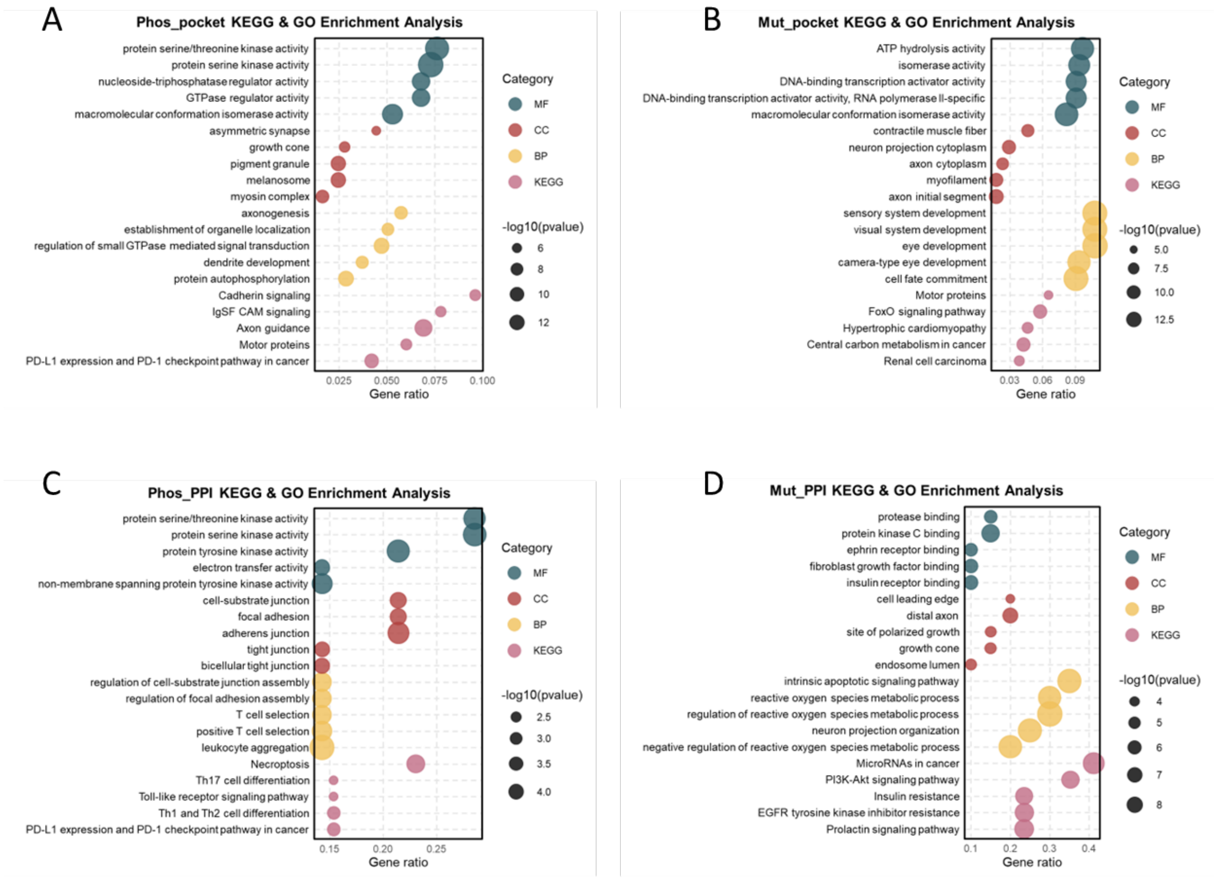


Figure S6. Functional enrichment analyses of proteins harboring structurally enriched disease-associated residues. (A) GO and KEGG enrichment of 852 proteins containing disease-associated phosphosites significantly enriched in ligand-binding pockets. (B) GO and KEGG enrichment of 639 proteins containing pathogenic mutations significantly enriched in ligand-binding pockets. (C) GO and KEGG enrichment of 15 proteins containing disease-associated phosphosites significantly enriched at PPI interfaces. (D) GO and KEGG enrichment of 20 proteins containing pathogenic mutations significantly enriched at PPI interfaces.

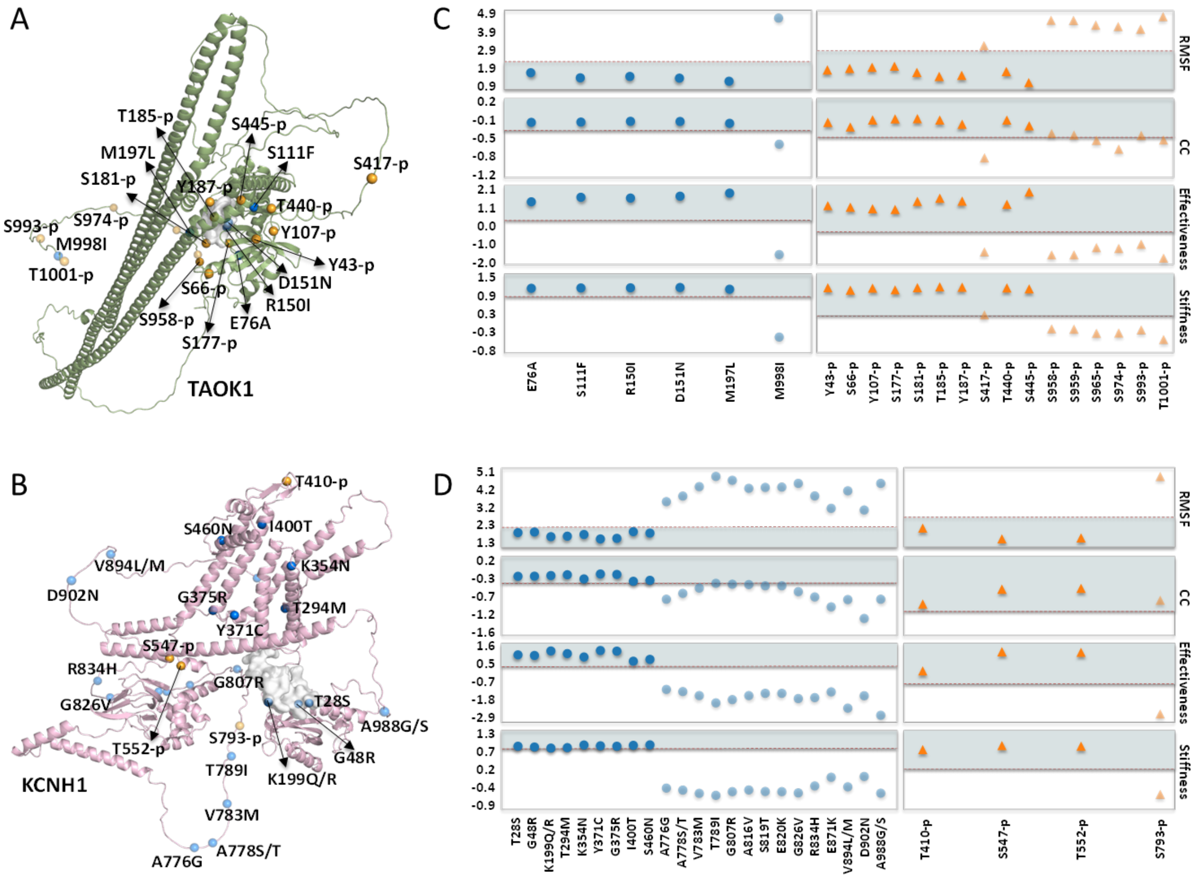

Figure S7. Supplementary case studies. (A, B) Structural representations and Fpocket-predicted pockets of the TAOK1 (A) and KCNH1 (B) systems. For both systems, only the single top-scoring Fpocket-predicted pocket overlapping with the positive sites is displayed. (C, D) Corresponding metric distribution plots for the TAOK1 (C) and KCNH1 (D) systems. All visualization, coloring, and transparency standards are consistent with those in Fig. 5.
