## Supplementary figures and images for "Dynamics-aware geometric learning predicts disease-associated molecular perturbations"

### Fig.S1.tif

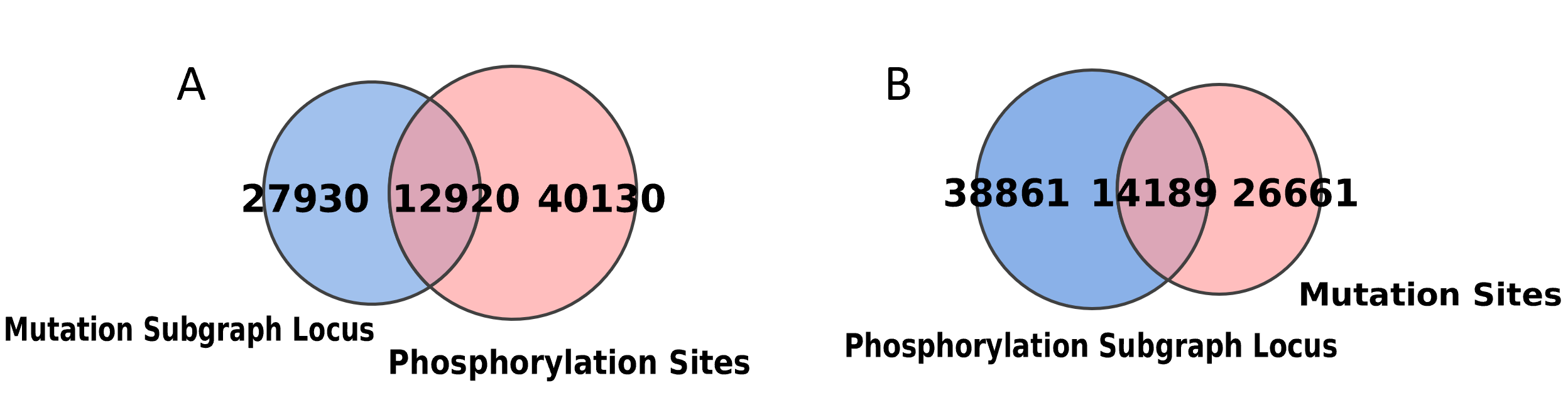

### Fig.S2.tif

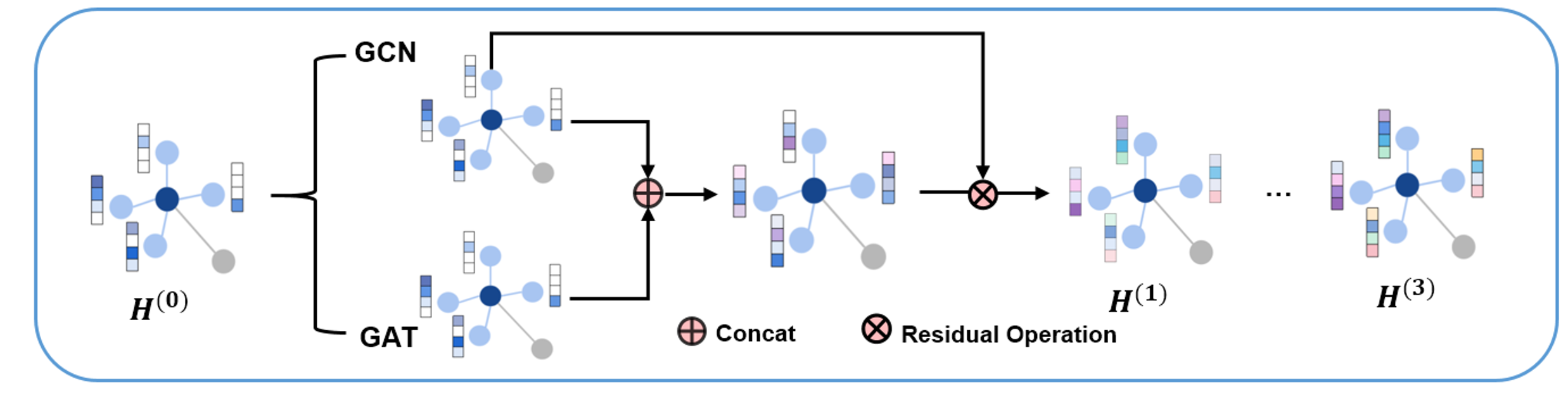

### Fig.S3.tif

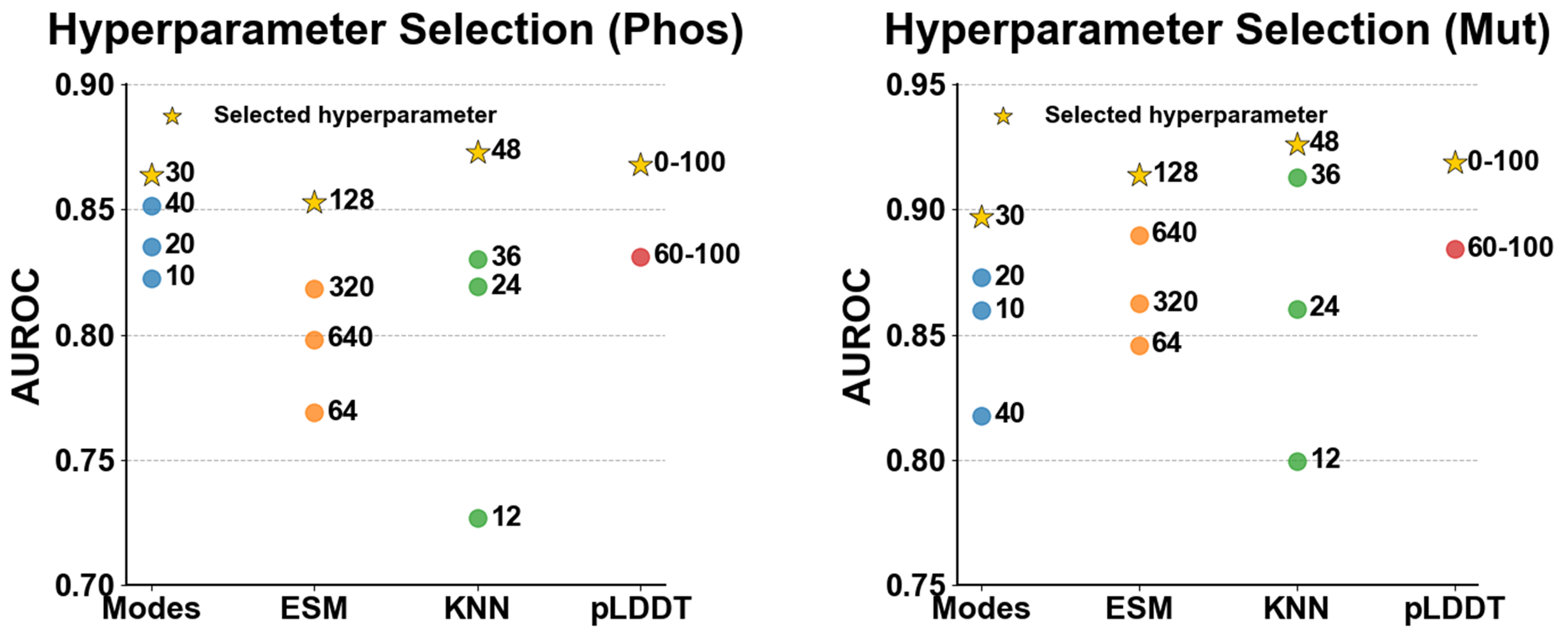

### Fig.S4.tif

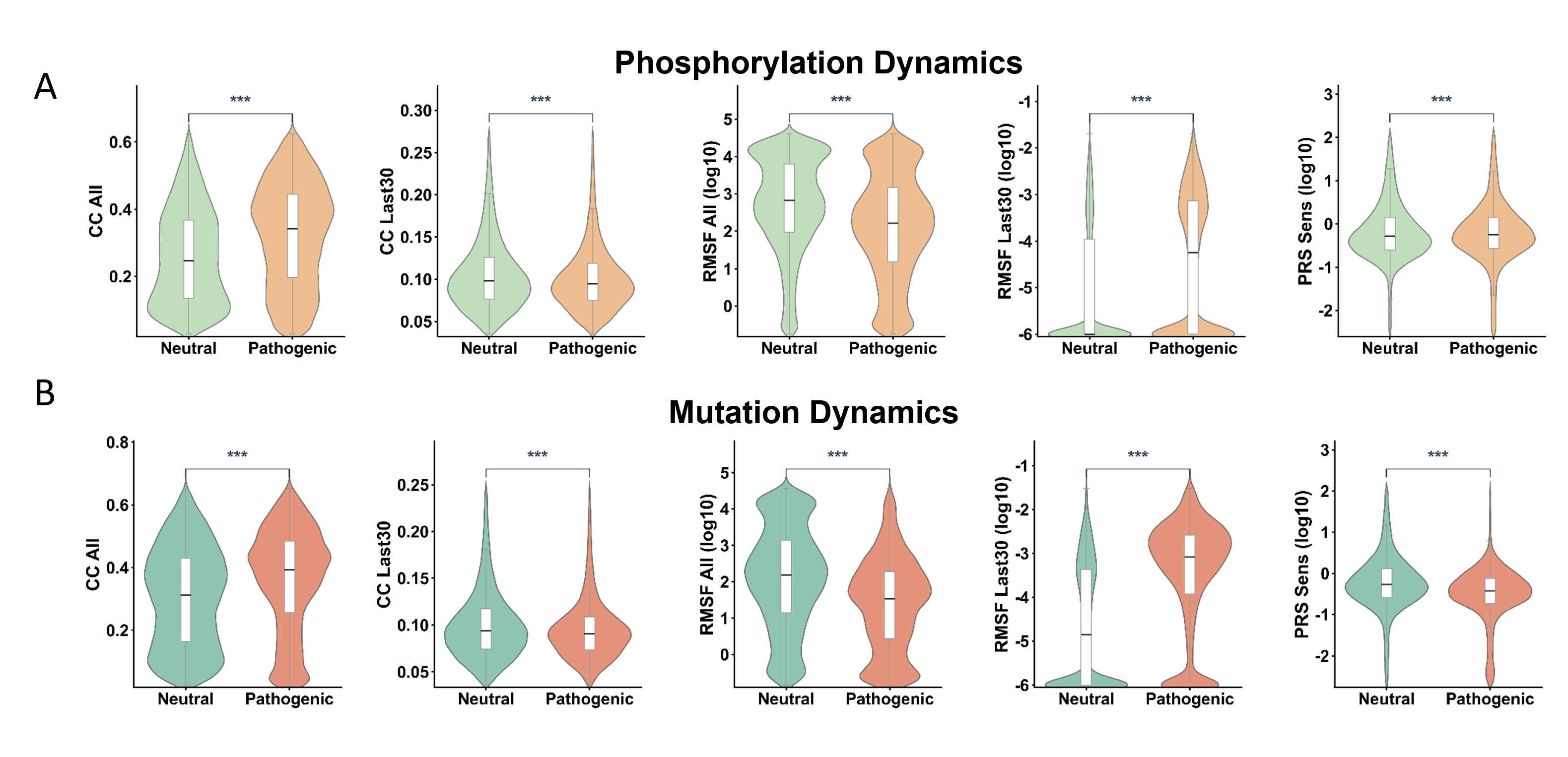

### Fig.S5.tif

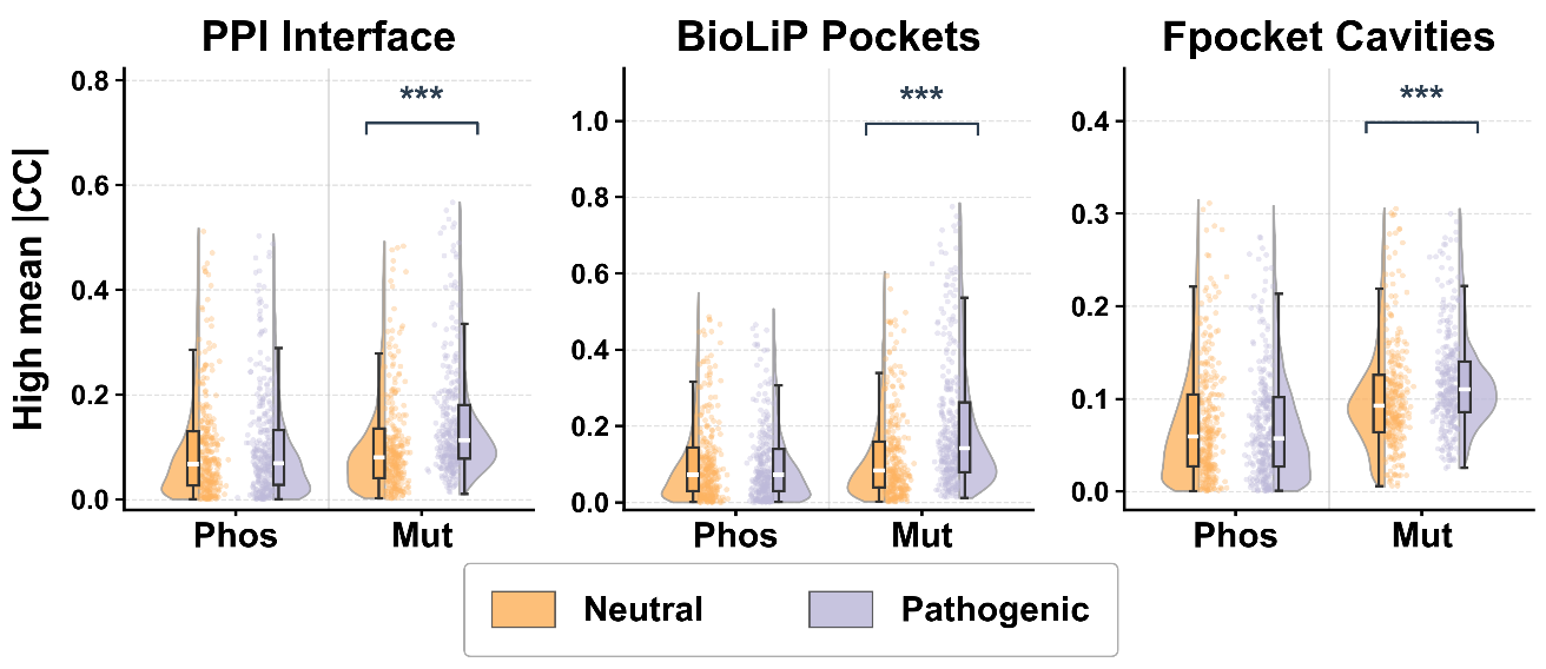

### Fig.S6.tif

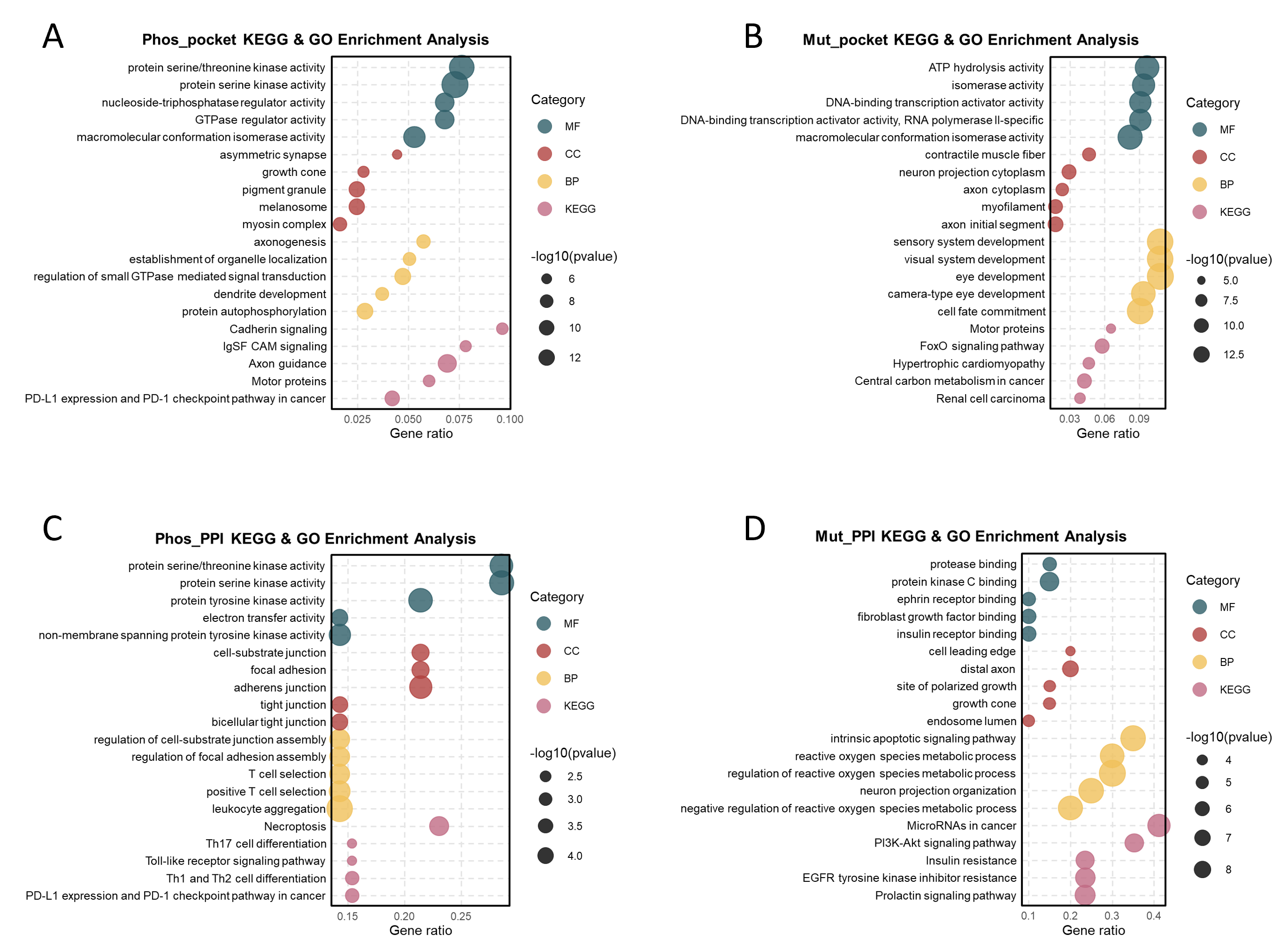

### Fig.S7.tif

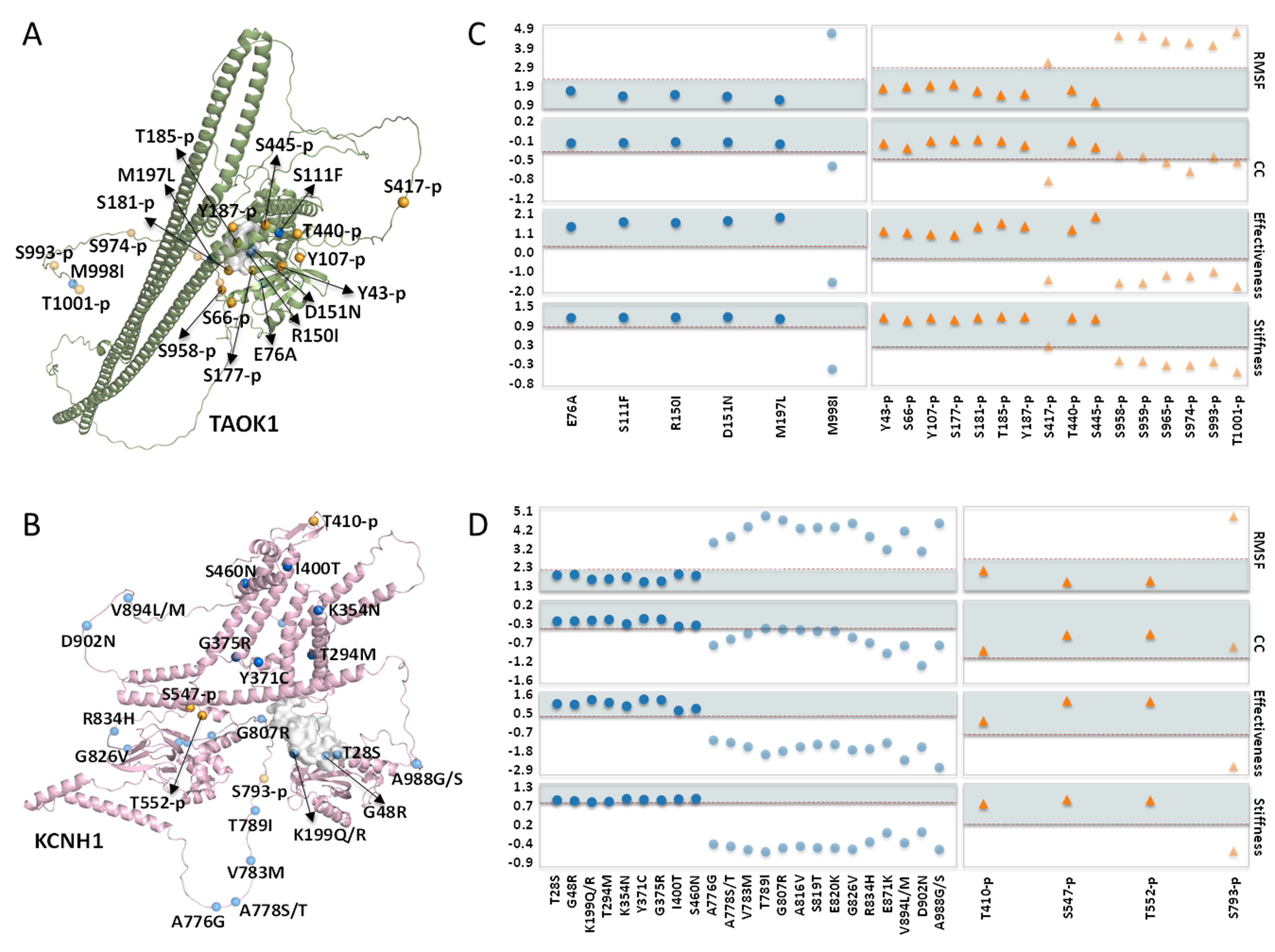
